# Real-time tracking of mRNP complex assembly reveals various mechanisms that synergistically enhance translation repression

**DOI:** 10.1101/2025.04.07.647595

**Authors:** Marco Payr, Julia Meyer, Eva-Maria Geissen, Janosch Hennig, Olivier Duss

**Affiliations:** Molecular Systems Biology Unit, European Molecular Biology Laboratory (EMBL), 69117 Heidelberg, Germany; Candidate for joint PhD degree from EMBL and Heidelberg University, Faculty of Biosciences, Heidelberg, Germany; Department of Biochemistry IV – Biophysical Chemistry, University of Bayreuth, 95447 Bayreuth, Germany; Data Science Centre, European Molecular Biology Laboratory (EMBL), 69117 Heidelberg, Germany

**Keywords:** Single-molecule FRET, translation regulation, RNA chaperone, protein-RNA interactions, mRNP complex formation, RNP dynamics

## Abstract

Protein biosynthesis must be highly regulated to ensure proper spatiotemporal gene expression and thus cellular viability. Translation is often modulated at the initiation stage by RNA binding proteins through either promotion or repression of ribosome recruitment to the mRNA. However, it largely remains unknown how the kinetics of mRNA ribonucleoprotein (mRNP) assembly on untranslated regions (UTRs) relates to its translation regulation activity. Using Sex-lethal (Sxl)-mediated translation repression of *msl-2* in female fly dosage compensation as a model system, we show that different mechanisms in mRNP assembly synergistically achieve tight translation repression. Using multi-color single-molecule fluorescence microscopy we show that 1) Sxl targets its binding sites via sliding and double-binding, 2) that Unr recruitment is accelerated over 500-fold by RNA-bound Sxl and 3) that Hrp48 further stabilizes RNA-bound Sxl indirectly via ATP-independent RNA remodeling. Overall, we provide a framework to study how multiple RBPs dynamically cooperate with RNA to achieve function.

## INTRODUCTION

Posttranscriptional regulation of gene expression is driven by essential mechanisms to precisely and spatiotemporally control protein biosynthesis (Buxbaum et al., 2015). This can be achieved via modulating RNA stability (Dave et al., 2023; Höpfler et al., 2023), localization (Mateju et al., 2020) or by regulating translation initiation (Lee et al., 2016). A misregulation of translation is linked to developmental defects and cancer (Siddiqui and Sonenberg, 2015; Tahmasebi et al., 2018). An important element in post-transcriptional gene regulation is the 3’ untranslated region (3’ UTR) which contains *cis*-regulatory elements that can determine the fate of the mRNA (Gebauer and Hentze, 2004). One example are microRNA induced silencing complexes that can bind to multiple target sites on the 3’ UTR to repress translation (Gebert and MacRae, 2019). Spatiotemporal control of translation is especially crucial to ensure correct organismal development, where for instance pre-mature *oskar* mRNA translation in *Drosophila* is inhibited by Bruno until the mRNA is localized to the posterior pole (Bose et al., 2022; Kim-Ha et al., 1995; Zimyanin et al., 2008). How multiple RNA binding proteins (RBPs) can act together on various regulatory RNA sequence elements and how the dynamics of mRNP complex assembly and disassembly contribute to efficient translation repression is not well-understood despite its relevance (He et al., 2023; Meyer et al., 2024).

To study how the kinetics of mRNA ribonucleoprotein (mRNP) complex assembly are linked to translation repression, we chose the Sex-lethal (Sxl)-mediated translation repression of *msl-2* in *Drosophila melanogaster* as a model system. Translational control of *msl-2* is needed for dosage compensation, which equalizes the expression of genes on the X chromosome between both sexes. Male flies require expression of MSL-2, a defining subunit of the dosage compensation complex (DCC), in order to achieve a two-fold increase in transcription of genes located on the X chromosome. In females, increased transcription of both X chromosomes would be lethal (Lucchesi and Kuroda, 2015). Multiple mechanisms have evolved to prevent expression of MSL-2 in females, one of which is translation repression of *msl-2* by the female-specific RBP Sxl (Bashaw and Baker, 1997; Kelley et al., 1997, p. 1997). On the *msl-2* mRNA, Sxl acts in a dual inhibitory way, where binding to poly(U) sites can block either ribosome scanning (5’ UTR) or pre-initiation complex recruitment (3’ UTR, Fig. 1A) and thus, ensures high fidelity in translation repression (Beckmann et al., 2005). The *msl-2* mRNA possesses three critical Sxl binding sites (B, E and F) out of 6 poly(U) tracts (A-F) on the *msl-2* transcript (Fig. 1A), but the necessity for having repeats for Sxl binding sites on the 3’ UTR (E and F) is not understood (Gebauer, 1999; Gebauer et al., 2003).

**Fig. 1.**
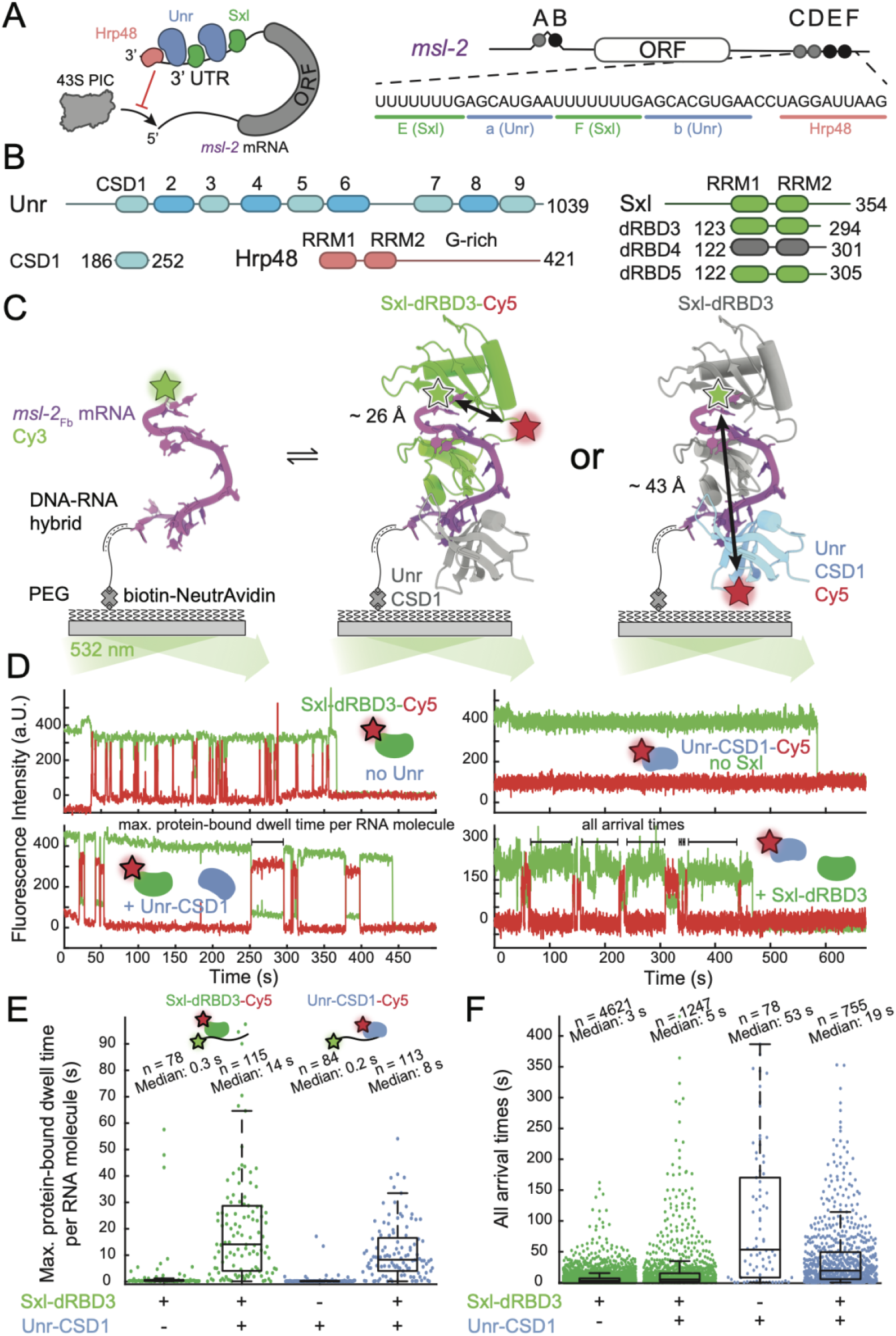
Sxl and Unr interact transiently with *msl-2* mRNA and stabilize each other. (A) Schematic depiction of the 3’ UTR-mediated translation repression of *msl-*2 by Sxl, Unr and Hrp48, which prevents 43S PIC recruitment. Binding sites of Sxl, Unr and Hrp48 within the 3’ UTR of *msl-*2 are shown in detail to the right. (B) Domain architecture and overview of constructs of Sxl, Unr and Hrp48 relevant to this study. (C) Structure-guided (PDB: 4QQB) experimental set-up to study the assembly of the Sxl-dRBD3 and Unr-CSD1 core repressive complex via single-molecule FRET. Labeling positions of either Sxl or Unr relative to the 5’ labeled immobilized RNA are indicated including distances between FRET pairs. (D) Representative single-molecule traces of labeled Sxl-dRBD3-Cy5 or Unr-CSD1-Cy5 (40 nM total protein) binding in absence of the other factor highlight the transient nature of binding. Addition of unlabeled Unr or Sxl prolongs the dwell times of the other factor. (E, F) Ensemble boxplots show the maximum protein-bound dwell times per RNA molecule (E) or all arrival times (F), respectively, with each dot representing a single protein binding event.

In addition to Sxl, Upstream of N-ras (Unr, Fig. 1A, 1B) is crucial for mediating translation repression. The *msl-2* transcript mediates cooperative Sxl and Unr protein-protein contacts, that lead to a high affinity ternary Sxl·Unr·RNA complex (Hennig et al., 2014). Mutations of specific Sxl residues, responsible for cooperative Sxl/Unr interactions, showed that a high affinity complex contributes to efficient translation repression, but it is unclear whether the high affinity arises from a faster recruitment of both factors to the *msl-2* mRNA or merely from a more stable, longer-lived, mRNP complex. This is especially important as it has been shown that both Sxl and Unr can bind their cognate binding sites *in vitro*, albeit with different affinities (Hennig et al., 2014). It raises the question whether Sxl and Unr bind stochastically to their cognate binding sites or if there is a sequential assembly with one protein getting recruited by another protein. Using a combination of mutational experiments and cross-linking studies (Duncan et al., 2006; Grskovic, 2003), it has been suggested that Unr gets recruited by Sxl, but direct molecular evidence in real-time for this proposed recruitment mechanism is missing.

Although the two RNA binding domains (RBDs) of Sxl, termed Sxl-dRBD3, are sufficient for protein-RNA complex formation, seven additional C-terminally located residues are critical for translation repression, termed Sxl-dRBD4 (Fig. 1B). However, a mechanistic understanding for the reduced translation repression activity of Sxl-dRBD3 is missing (Grskovic, 2003).

Furthermore, the 3’ UTR-mediated translational repression requires the factor Hrp48 (Szostak et al., 2018). The Hrp48-RRM12 RNA binding motif on the *msl-2* transcript (Lomoschitz et al., 2025) has been mapped and its relevance in translation repression of *msl-2* has been demonstrated (Szostak et al., 2018) (Figure 1A, B). However, it remains unclear whether Hrp48 establishes direct contacts to Unr or Sxl and by which mechanisms it contributes to translation repression.

Here, we combine multi-color single-molecule fluorescence microscopy assays with nuclear magnetic resonance spectroscopy (NMR) and reporter gene assays to study the mechanism for efficient translation repression of the *msl-2* mRNA by a dynamic multi-component mRNP complex assembly. Simultaneously monitoring the formation and dissociation of multiple protein-RNA interactions in real-time via single-molecule Förster resonance energy transfer (smFRET), we show that mRNP complex assembly on the *msl-2* transcript is highly dynamic and relies on multiple synergistic effects required for efficient translation repression. The repressive Sxl-mRNA interaction is increased via Sxl sliding on mRNA or can be enhanced by the binding of a second Sxl protein. RNA-coupled protein folding of the intrinsically disordered C-terminal extension of Sxl into an α-helix enhances its translation repression activity. Furthermore, a cooperative RNA-mediated complex assembly between Sxl and Unr, gives rise to a 500-fold increase in recruitment rates of Unr to the *msl-2* transcript. Finally, we show that Hrp48 indirectly contributes to Sxl·mRNA complex stability by RNA remodeling rather than by direct protein-protein interactions, explaining its contribution to translation repression activity. Overall, our work provides a quantitative biophysical understanding of how several multi-domain eukaryotic RNA binding proteins dynamically cooperate on a common 3’ UTR using different mechanisms to efficiently repress translation initiation.

## RESULTS

### Ternary Sxl·Unr·*msl-2* complex formation is transient and reversible

In order to detect the binding dynamics of Sxl and Unr to the *msl-2* mRNA 3’ UTR in real-time, we immobilized the target RNA, fluorescently labeled with a Cy3 dye, to a glass slide and added site-specifically Cy5-labeled proteins to the reaction solution for single-molecule imaging. In a previously published crystal structure (Hennig et al., 2014, PDB: 4QQB), the RNA-binding domains of Sxl and Unr, Sxl-dRBD3 (*Drosophila* RNA binding domains construct 3, residues 123-294) and Unr-CSD1 (cold shock domain 1, residues 185-252, Fig. 1B), were used to determine the RNA-binding interface and protein-protein contacts of Sxl and Unr to site F in the 3’ UTR of the *msl-2* mRNA (Fig. 1A). Based on the structure, we engineered single-cysteine mutants of Sxl-dRBD3 (C154S, S207C) and Unr-CSD1 (C23S) for site-specific fluorophore tagging. We placed the cysteines such that fluorescent labeling minimally interferes with the RNA binding interface (Fig. 1C) and ensured that the distance between the acceptor dye on the protein and the donor dye on the target RNA results in a high FRET signal (Sxl-mRNA distance ∼ 26 Å; Unr-mRNA distance ∼ 43 Å). We first studied the binding of both proteins to the *msl-*2 mRNA, where we used an mRNA that contains the Sxl binding region F (U_7_G tract) together with its downstream Unr binding region, termed region b in this study (Fig. 1A), because this sequence (construct called *msl-2*_Fb_) has the strongest effect on translation repression of the *msl-2* mRNA (Gebauer et al., 2003). Monitoring the binding of Sxl-dRBD3-Cy5 to *msl-2*_Fb_ in real-time showed that bound dwell times are very short with a median maximum dwell time per RNA molecule of 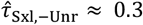 s (Fig. 1D, E). In comparison, the addition of unlabeled Unr-CSD1 increased the median maximum dwell time per RNA molecule of Sxl by almost 50-fold (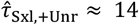 s). The increase in Sxl-bound dwell times in presence of Unr is in agreement with protein-protein contacts of Sxl and Unr in the ternary Sxl·Unr·*msl-2* complex structure (Hennig et al., 2014, PDB: 4QQB). The addition of Unr-CSD1 also showed that all Sxl-bound dwell times switch from a single-exponential to a double-exponential distribution (Figure S1A). The mixture of populations can be described by a bound lifetime of longer (⟨τ⟩ = 8.8 ± 0.3 s; 38 ± 1 %) and shorter duration (⟨τ⟩ = 0.49 ± 0.03 s; 62 ± 1 %) (Fig. S1A, S1B). We attribute the short and long bound lifetimes to Sxl binding in absence or presence of Unr co-bound to the same RNA molecule, respectively.

Having investigated the binding dynamics of Sxl, we next investigated the binding kinetics of Unr by monitoring binding of Unr-CSD1-Cy5 to the *msl-2*_Fb_ mRNA. We observed very few and short-lived observable dwell times (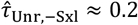s, Fig. 1D, E). Upon addition of unlabeled Sxl-dRBD3, the median maximum Unr dwell time per RNA molecule increased by a factor of roughly 40 to 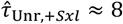 s (Fig. 1E). However, not only the RNA-bound dwell time of Unr is affected by the presence of Sxl - also the median arrival time 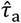 of observable Unr binding events is decreased (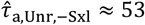 s; 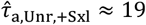s; [Unr-CSD1-Cy5] = 26 nM) (Fig. 1F). Overall, our single-molecule data support the cooperative binding relationship between Sxl and Unr found in previous structural studies (Hennig et al., 2014) and provide a first understanding of the real-time binding dynamics of these two proteins to RNA.

### Non-canonical Sxl interactions with the *msl-2* mRNA contribute to translation repression

Dwell times of both Unr and Sxl were surprisingly short and we therefore wanted to understand if an increase in mRNP complex dwell time can act as a hallmark of translation repression. For translation repression of *msl-*2, it is known that the RNA-binding domains of Sxl, Sxl-dRBD3, although sufficient for complex formation *in vitro*, have a significantly weaker translation repression activity compared to a construct, which is C-terminally extended by seven amino acids (Sxl-dRBD4, residues 122-301, Figure 1B) (Gebauer, 1999; Grskovic, 2003). However, it is unclear as to why. AlphaFold2 predicts an α-helix for a conserved region C-terminal of the RNA binding domains. An overlay of the structure of the ternary complex of Sxl-dRBD3·Unr-CSD1·*msl-2* (Hennig et al., 2014; Jumper et al., 2021) with the complex containing the C-terminally predicted α-helix shows no steric clashes (Fig. 2A, S2A, S2B). Based on sequence conservation and the structural prediction we designed a new Sxl construct, termed Sxl-dRBD5 (122-305, 51B), that is 11 amino acids C-terminally extended compared to Sxl-dRBD3.

**Fig. 2.**
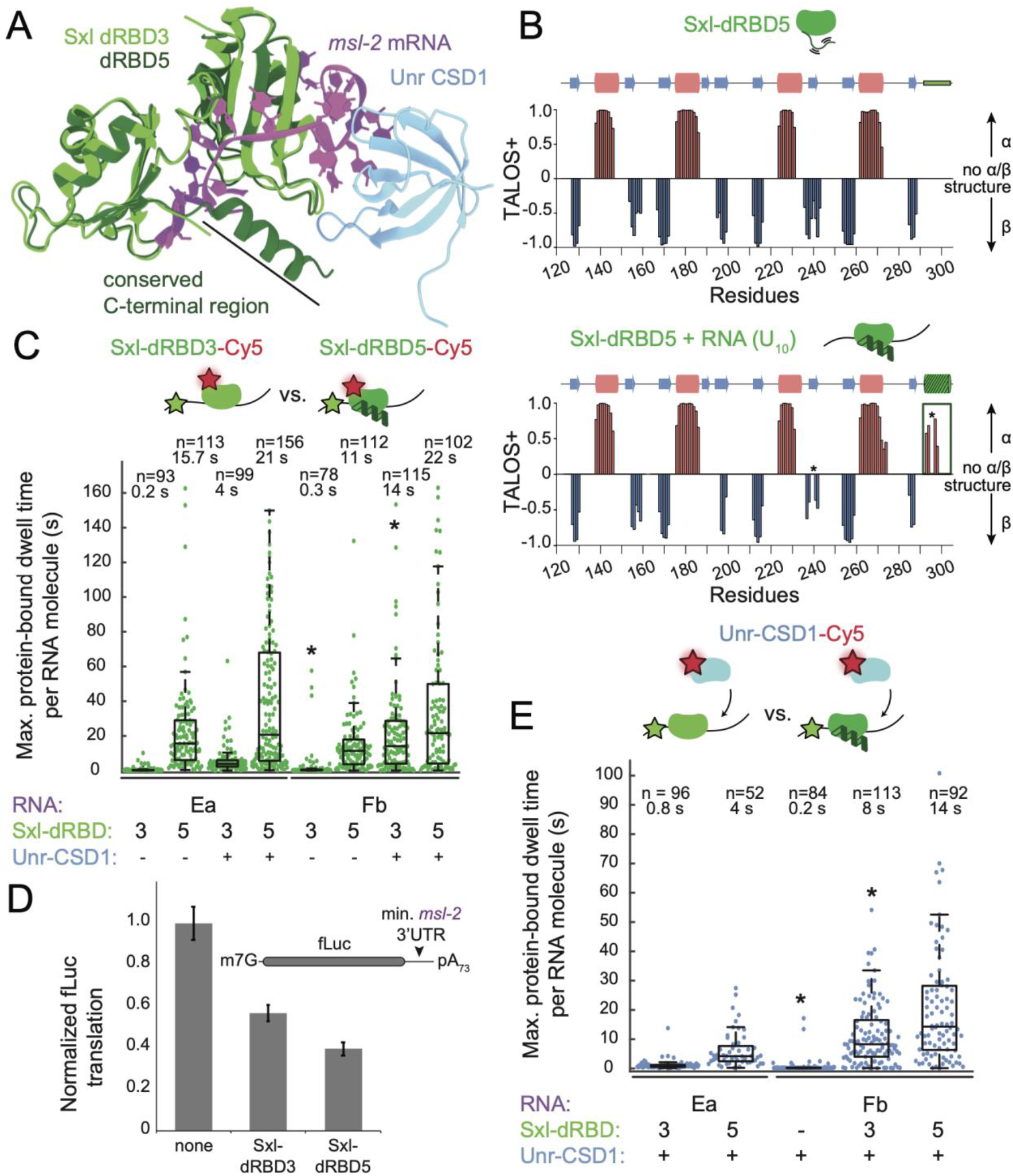
Sxl participates in non-canonical RNA interactions to aid translation repression. (A) Overlay of an AlphaFold2 prediction of Sxl-dRBD5 aligned to Sxl-dRBD3 in a ternary complex with Unr and *msl-2*_Fb_. (B) TALOS+ secondary structure predictions based on backbone chemical shifts of Sxl-dRBD5 show that C-terminal residues are unfolded in the absence of RNA. Addition of U_10_-mer RNA increases the helical propensity of C-terminal residues. Asterisks denote unassigned residues. (C) Binding of labeled Sxl constructs (Sxl-dRBD3-Cy5 compared to Sxl-dRBD5-Cy5) is shown for different binding sites (*msl-2*_Ea_ and *mls-2*_Fb_) in absence and presence of unlabeled Unr-CSD1. (D) Translation repression activity is increased in presence of Sxl-dRBD5 in comparison to Sxl-dRBD3 (1.2 µM each) as measured using a reporter mRNA that contains the minimal 3’ UTR of *msl-*2. (E) Binding of Unr-CSD1-Cy5 to both binding sites (*msl-2*_Ea_ and *mls-2*_Fb_) with and without unlabeled Sxl-dRBD3 or Sxl-dRBD5. Experiments that are marked with an asterisk are also shown in Fig. 1E.

To confirm the presence of the predicted C-terminal helix, we utilized NMR to detect secondary structure changes of Sxl-dRBD5 by measuring secondary chemical shifts in absence and presence of RNA. This revealed that the C-terminal region is dynamic and transitions from being intrinsically disordered in the apo state to an α-helical conformation in the RNA bound state (Fig. 2B, S2C). While we observed chemical shift perturbations (CSPs) in this C-terminal region of Sxl-dRBD5 in NMR experiments upon titration of its cognate RNA motif, no additional CSPs are observed upon addition of Unr-CSD1 to the RNA-bound Sxl-dRBD5 (Fig. S2D, S2E), which suggests that there are no additional interactions between Sxl-dRBD5 and Unr-CSD1 in presence of this additional C-terminal α-helix.

Next, to investigate the functional effect of the additionally identified transient C-terminal α-helix in Sxl, we performed fluorescence polarization assays (Fig. S3A, S3B) and ITC measurements (Fig. S4A-C) which show that the C-terminal helix increases affinity of Sxl towards its binding site. Furthermore, we compared the RNA binding dynamics of Sxl-dRBD5-Cy5 and Sxl-dRBD3-Cy5 using smFRET. The C-terminal extension of Sxl results in a median maximum Sxl-bound dwell time per molecule which is increased by 35-fold for the *msl-2*_Fb_ construct (Fig. 2C) and reduces observable arrival times 5-fold for both E and F sites (Fig. S4D). Finally, we tested the functional relevance of this transient C-terminal α-helix, using *in vitro* translation assays with an mRNA that carries a firefly luciferase gene and the minimal 3’ UTR of *msl-2*. We find that Sxl-dRBD5 has a 17 ± 5 % increased translation repression activity compared to Sxl-dRBD3 (Fig. 2D, S3C). Together, our structural, biophysical and functional data demonstrate that a short amino-acid extension at the C-terminus of the well-characterized RNA binding domain of Sxl forms a transient α-helix which is needed for full translation repression (Gebauer et al., 2003) and that the increased Sxl bound dwell time to its target RNA correlates with translation repression activity.

### Multi-color smFRET reveals hierarchy in assembly and vastly accelerated recruitment of Sxl and Unr in presence of the other protein

It was previously proposed that Sxl recruits Unr (Duncan et al., 2006; Grskovic, 2003) but experimental evidence is missing. While fluorescently labeling one protein at a time allowed us to conclude that both Sxl and Unr increase the RNA-bound dwell times of each other, it is not clear which protein initiates the ternary complex assembly. In order to simultaneously detect the binding of both proteins to a single RNA target molecule (*msl-2*_Fb_ mRNA), we set up a three-color single-molecule FRET assay, by adding Sxl-dRBD5-Cy5 and Unr-CSD1-Cy5.5 to the reaction. Our extended assay allows detection of three different binding modes (Fig. 3A): 1) isolated Sxl (Cy3-Cy5 FRET), 2) isolated Unr (Cy3-Cy5.5 FRET), and 3) co-binding events (Cy3-Cy5 & Cy3-Cy5.5 FRET). We could observe all three binding modes in this assay (Fig. 3B, Fig. S6). The vast majority of co-binding events showed first Sxl-dRBD5-Cy5 binding (83 %), which was followed by multiple re-binding events of Unr during a single long-lived Sxl binding event (Fig. 3C). Only a small fraction of co-binding events (4 %) showed that Unr-CSD1-Cy5.5 bound first. The distribution of binding modes suggests that the assembly of the ternary Sxl·Unr·*msl-2* complex follows a hierarchical order, in which stable Unr binding requires initial recruitment of Sxl. In 13 % of the Sxl/Unr co-binding events, we could not determine which protein arrived first, because the assembly took place in less than 200 ms (limited by our experimental frame rate). Such a high fraction of fast assemblies indicates that Sxl/Unr mRNP complex formation is extremely fast. Indeed, the median number of binding events per second 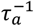 show that Unr recruitment by Sxl is increased by approximately 500-fold (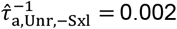 s^-1^, 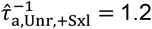 s^-1^). Vice versa, in the rare case of Unr binding first, Sxl recruitment is accelerated by 560-fold (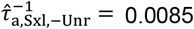 *s*^−1^, 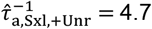 s^-1^, Fig. 3D). The hierarchy in assembly and the accelerated recruitment of both factors was also observed with other protein labeling schemes (Fig. S5). Overall, our multi-color smFRET assays demonstrate that Sxl recruits Unr and boosts its recruitment rate by more than two orders of magnitude highlighting how cooperativity can rely both on off- and on-rates to ensure efficient RNP assembly.

**Fig. 3.**
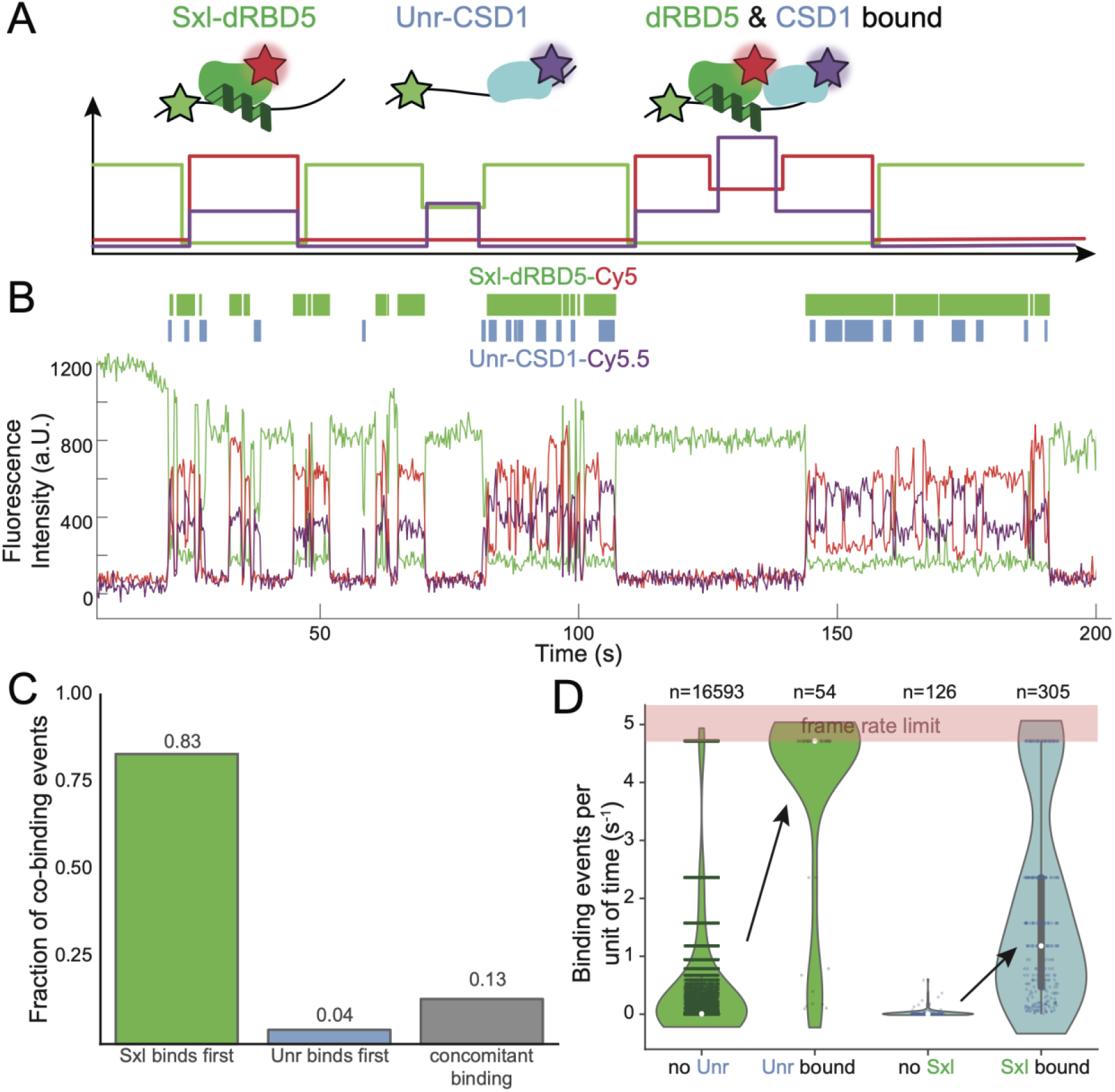
A hierarchical Sxl and Unr assembly is vastly accelerated in the presence of the other factor. (A) Schematic depiction of the three possible assembly scenarios that can be observed: 1) bound Sxl in isolation, 2) bound Unr in isolation and 3) both proteins co-bound to the mRNA. (B) Representative trace of a three-color smFRET experiment to study multi-component mRNP assembly with overlapping Sxl and Unr dwell times. Concentrations: Sxl-dRBD5-Cy5 (40 nM total) and Unr-CSD1-Cy5.5 (200 nM total). See Fig. S6 for full trace. (C) Relative distribution of the order of assembly for co-binding events. (D) Binding events per second for labeled Sxl (green) and Unr (blue) were derived from co-binding events where Sxl and Unr binding overlapped. Binding events per second for Unr-CSD1-Cy5.5 (200 nM total) in absence of Sxl and binding events per second for Sxl-dRBD5-Cy5 (40 nM total) in absence of Unr were derived from different experiments.

### Sxl-mRNA association time increases by sliding and simultaneous binding of two Sxl copies

For efficient translation repression of the *msl-2* mRNA, two repeats of U-rich tracts, the E and F sites, are required (Gebauer et al., 2003). Having investigated the Sxl binding dynamics to isolated *msl-2*_Ea_ and *msl-2*_Fb_-sites so far, we next extended our single-molecule assays to study the binding kinetics of Sxl to an immobilized RNA construct encompassing both E and F sites simultaneously, referred to as *msl-2*_EaFb_ (Fig. 4A, B). The median of all Sxl-bound dwell times increases for the tandem *msl-2*_EaFb_ construct 30-fold (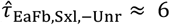 s) compared to the isolated *msl-2*_Ea_ and *msl-2*_Fb_-sites (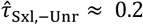 s for *msl-2*_Ea_ and *msl-2*_Fb_) (Fig 4C). The binding events of Sxl-dRBD5-Cy5 to *msl-2*_EaFb_ revealed states with high (⟨*E*_*HF*_⟩ ≈ 0.9) and low FRET (⟨*E*_*LF*_⟩ ≈ 0.3) efficiency, that we assign to E and F site binding, respectively (Fig. 4B, D). We confirmed this assignment using an E-site mutant (*msl-2*_EmaFb_) showing mostly low FRET (Fig. S7D). However, high and low FRET binding events did not only occur in isolation, but also interchanged throughout a single binding event of Sxl suggesting sliding of a single Sxl protein molecule from one binding site to the other (Fig. 4D, S7A).

**Fig. 4.**
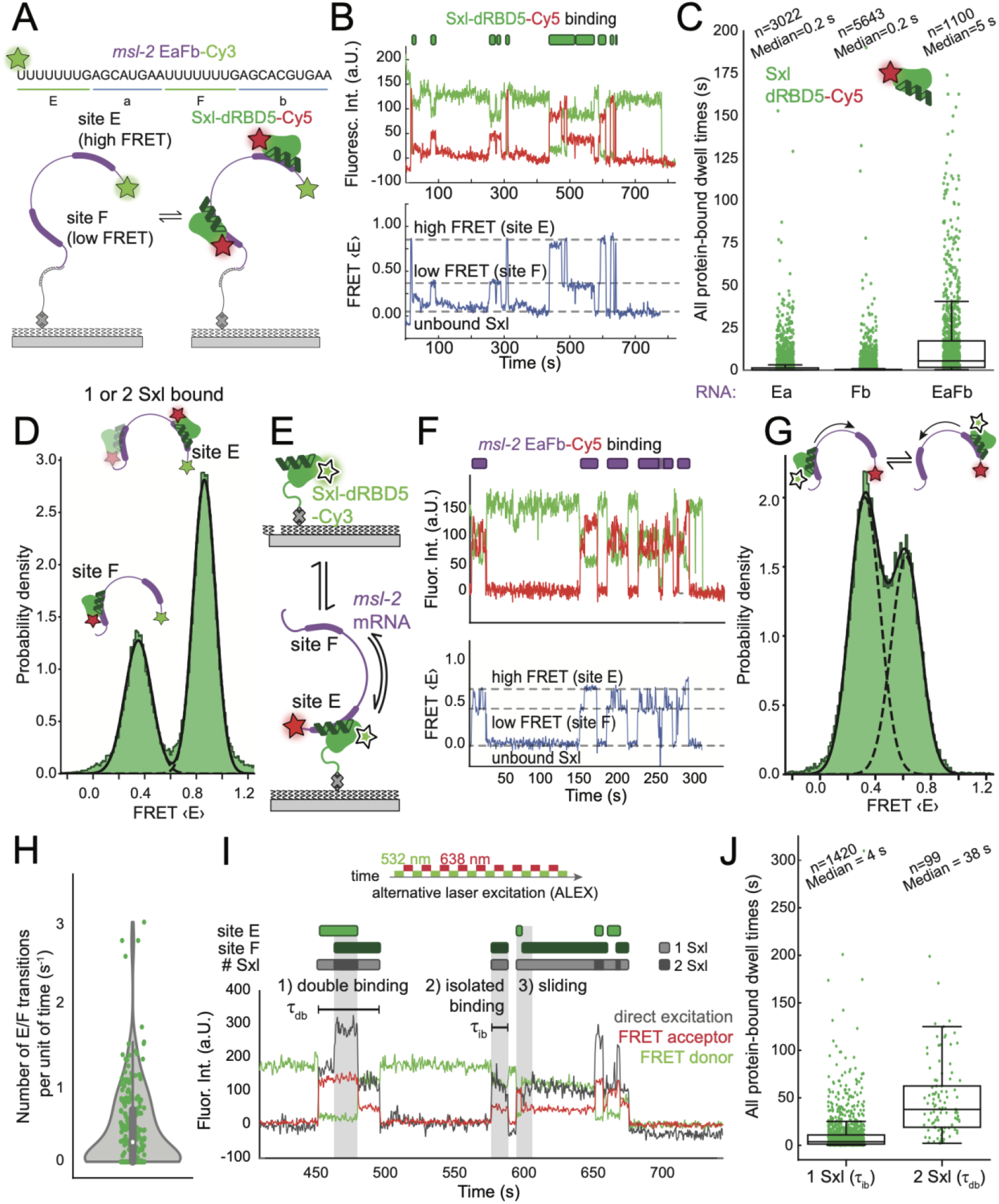
Sxl can slide along the *msl-2* mRNA and bind two sites simultaneously. (A) Schematic depicition of Sxl-dRBD5-Cy5 binding experiments to a tandem RNA construct including E and F sites. (B) Representative trace of Sxl binding to the *msl-2*_EaFb_ mRNA shows both high-FRET and low-FRET binding events marked with green boxes. (C) A comparison of all Sxl-bound dwell times to isolated *msl-2*_Ea_, *msl-2*_Fb_ RNA and tandem *msl-2*_EaFb_ RNA. (D) 1D FRET histogram of binding events shows that both E and F sites can be bound. The histogram was fitted to a linear combination of two gaussians. (E) Illustration of a 1D sliding assay in which single-immobilized AviTag-Sxl-dRBD5-Cy3 is transiently bound by *msl-2*_EaFb_-Cy5 mRNA. (F) Binding of *msl-2*_EaFb_ mRNA to single-Sxl molecules shows both high-FRET and low-FRET binding modes, representing binding to either E and/or F-sites, respectively. (G) 1D histogram of the sliding assay. (H) Frequency of FRET state transitions was evaluated for each binding event. (I) Exemplary trace of Sxl-dRBD5-Cy5 binding in an ALEX experiment (100 ms exposure time per laser) demonstrates that Sxl can bind both sites 1) simultenously, 2) in isolation, or 3) a single Sxl protein can slide between both E and F sites. (J) Quantification of isolated and double Sxl binding events shows that double Sxl occupancy leads to a 10-fold longer total Sxl-bound dwell time.

To further investigate if sliding can contribute to longer bound dwell times of Sxl on *msl-2*_EaFb_, we designed a sliding FRET assay by immobilizing a single Sxl-dRBD5-Cy3 protein to the imaging surface and adding *msl-2*_EaFb_-Cy5 RNA to the flow chamber (Fig. 4E). We observed again two interchanging FRET states during single binding events for 61 % of the events (Fig. S7B), further supporting our model that Sxl slides between both E and F binding sites on the *msl-2*_EaFb_ 3’ UTR (Fig. 4F, 4G). The transition frequency of Sxl between both E and F-sites occurs with a median frequency of 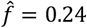 s^-1^ (Fig. 4H). This is on a similar timescale as Unr recruitment by Sxl (Fig. 3D, 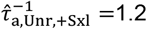 s^-1^), which suggests that both Sxl sliding between E and F sites and fast Unr recruitment can occur with a similar probability. The distribution of interchanging FRET states revealed that this movement is highly dynamic and heterogenous between binding events, where a substantial fraction of binding events does not exhibit any switch in FRET states (39 %) and fully resides in either low or high FRET state during binding (Fig. S7B). We also reason that the exchange of FRET states cannot be due to conformational dynamics of the RNA as a mutation of the E-site almost completely abolishes the high FRET states (Fig. S7D).

Having established that a single Sxl protein can remain associated longer to its target RNA by having the possibility to diffuse to the other binding site instead of immediately dissociating, we wondered whether two Sxl protein molecules could simultaneously bind to the *msl-2*_EaFb_-tandem construct. To this end, we immobilized *msl-2*_EaFb_-Cy3 to the imaging surface and added Sxl-dRBD5-Cy5 to the reaction, imaging with alternative laser excitation (ALEX). This experimental setup allows us to simultaneously detect which of the two sites (E or F) are occupied by a Sxl protein and also informs on the protein stoichiometry (Fig. 4I). Our data show that in addition to sliding, also both binding sites can be occupied simultaneously by Sxl at physiological concentrations. We quantified the median of all Sxl-bound dwell times for Sxl bound in isolation or when co-bound with another Sxl protein and find an almost 10-fold increase in total Sxl bound-dwell time in presence of two Sxl protein copies on the same mRNA molecule (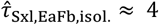 s; 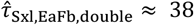 s) (Fig. 4J). Overall, our data demonstate that the tandem *msl-2*_EaFb_ sites increase Sxl-RNA bound dwell time by both sliding of a single Sxl protein between both sites or by occupancy of both sites by two Sxl proteins (Fig. 4I). This complex target recognition mode of Sxl functionally leads to increased translation repression efficiency of the *msl-2* mRNA (Gebauer et al., 2003).

### Hrp48 increases Sxl-bound dwell time indirectly by resolving inhibitory RNA structures

It has been shown that Sxl and Unr alone are not sufficient for 3’ UTR-mediated translation repression of the *msl-2* mRNA (Szostak et al., 2018). Having investigated the binding kinetics of Sxl and Unr, we next biophysically characterized another factor that was previously shown to be important for translation repression of the *msl-2* mRNA. Hrp48 has been suggested to be involved in 3’ UTR-mediated translation repression, but the interdependency of Sxl/Unr with Hrp48 remains unanswered (Szostak et al., 2018). To this end, we repeated our single-molecule assays by immobilizing a 3’ UTR construct comprising the EF-site and the entire downstream sequence, including the Hrp48 binding site and poly(A) tail (Fig. 5A) (Lomoschitz et al., 2025). Because we already observed very stable binding of Sxl-dRBD5 in the presence of Unr-CSD1 in our single-molecule assays (Fig. 2C), we increased the ionic strength from 50 mM to 150 mM and the temperature from 21 °C to 29 °C. This decreased the Sxl-bound dwell times in presence of Unr (maximum Sxl-bound dwell time per RNA molecule: 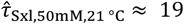 s & 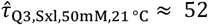s; 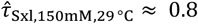 s & 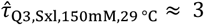 s, Fig. S8) enabling us to detect stabilizing effects of other RBPs without being limited by photobleaching. With these more physiological experimental parameters, the median and 75^th^ percentile (Q3) maximal dwell time of Sxl-dRBD5-Cy5 binding to the 3’ UTR was substantially increased by the addition of both full-length Unr and Hrp48 (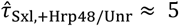 s; 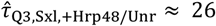 s), but less substantial by the addition of Unr (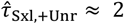 s; 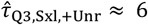 s) or Hrp48 alone (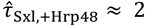 s; 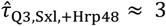 s)(Fig. S9B). Similarly, we observed that Hrp48 can independently bind to *msl-2* and that addition of Sxl/Unr stabilizes bound dwell times of Hrp48 (Fig. S9D, S9E), further supporting a cooperative binding relationship between Sxl/Unr and Hrp48.

**Fig. 5.**
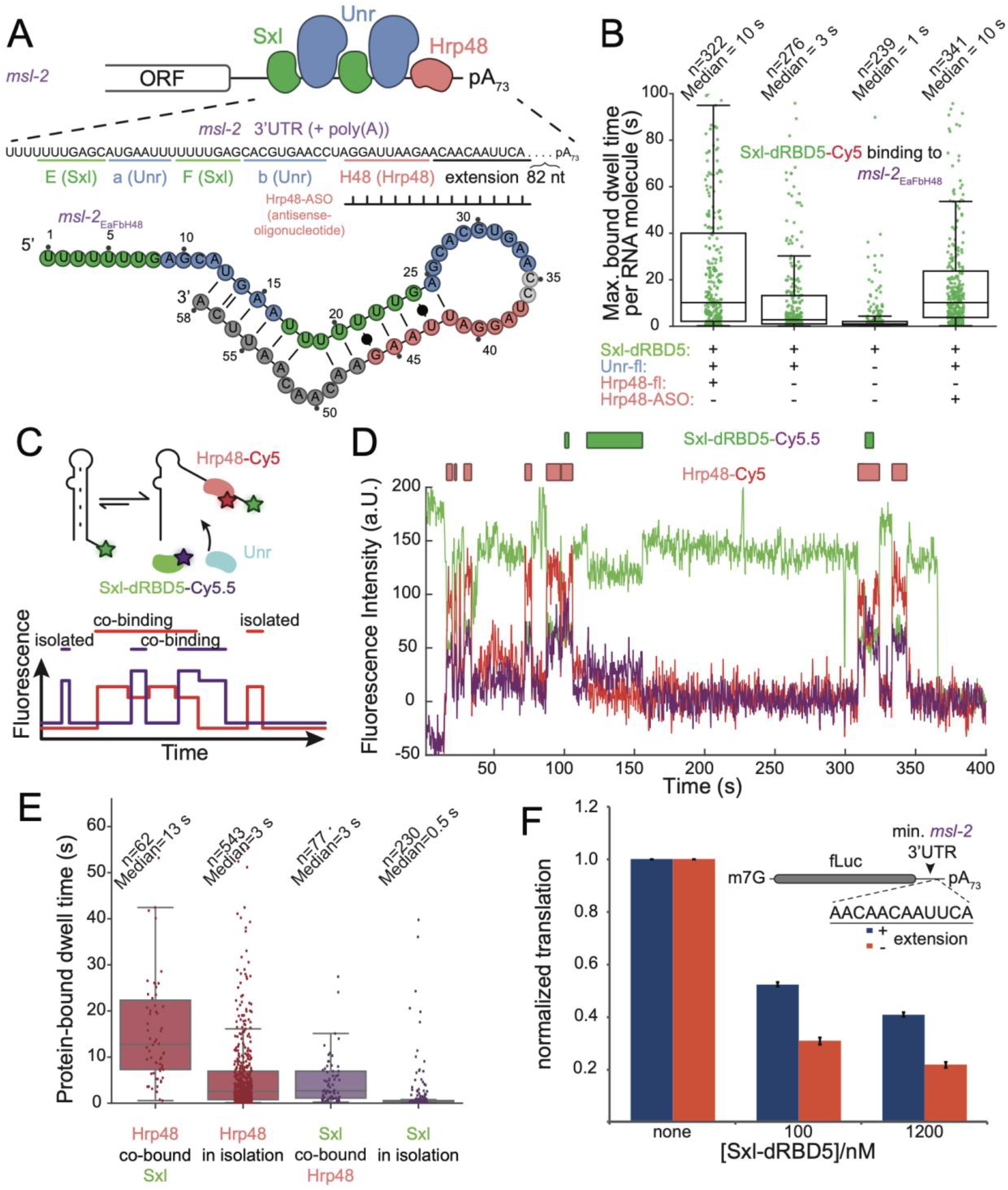
Hrp48 chaperones the *msl-*2 3’ UTR to maintain stable Sxl·Unr·RNA association. (A) Schematic depiction of RBPs that are involved in 3’ UTR-mediated translation repression of the *msl-2* mRNA. RNA constructs used in smFRET assays are shown below together with an RNA secondary structure prediction of *msl-2*_EaFbH48_ (Zuker, 2003). Hrp48-ASO describes the antisense oligonucleotide that targets the Hrp48 binding site including the 3’ extension. The 3’ UTR RNA construct starts at the first Sxl binding site E and comprises the full downstream sequence including a poly(A)_73_ tail. (B) Sxl-dRBD5-Cy5 binding to a shorter RNA (*msl-2*_EaFbH48_) that comprises binding sites for Sxl, Unr and Hrp48. Addition of an antisense oligonucleotide (Hrp48-ASO) that targets the Hrp48 binding site including an extension rescues the Sxl stabilizing effect. Experiments were done in duplicates. Individual replicates are shown in Fig. S9. (C) Scheme of a multicolor experiment to simultaneously track Hrp48 and Sxl binding. (D) A representative experimental trace shows co-bound Hrp48-Cy5 and Sxl-dRBD5-Cy5.5 events. (E) Ensemble distribution of all dwell times shows that co-bound dwell times of the individual labeled proteins are longer compared to isolated binding events. (F) Bulk translation assays show that an mRNA reporter lacking the repressive 3’ extension, which base-pairs with the E/F-site, is repressed more efficiently by Sxl.

With this assay we were able to capture a previously unobserved link between Hrp48 and the Sxl/Unr translation repression proteins. This was unexpected because previous studies could not observe a direct interaction between the RNA-binding domains of Hrp48 and Sxl in absence or presence of RNA (Lomoschitz et al., 2025; Szostak et al., 2018). Using mass photometry, we also did not observe an interaction between full-length Hrp48 and Sxl-dRBD5 or Hrp48 and full-length Unr (Fig. S10A, S10B). We therefore hypothesized that Hrp48 may exert its effect on Sxl binding indirectly via RNA remodeling, rather than by direct Hrp48-Sxl/Unr protein-protein interactions.

To investigate this possibility, we created a simplified RNA construct consisting of the EaFb site, the downstream Hrp48 binding site and a short 12 nucleotides native 3’-sequence extension (*msl-2*_EaFb-H48_). The shortening of the *msl-2* mRNA construct reproduced the increased Sxl-bound dwell times in presence of Hrp48 and Unr (Fig. 5B, S9B, S9C). The predicted RNA secondary structure (Zuker, 2003) of this minimal construct is in agreement with a model in which the binding sites of Sxl and Unr are sequestered by intramolecular interactions with the Hrp48 binding sites and its downstream sequence extension (Fig. 5A). We hypothesized that binding of Hrp48 would weaken RNA secondary structure or prevent its formation. In order to test this hypothesis, we substituted the Hrp48 protein with an antisense DNA oligonucleotide (Hrp48-ASO) that can anneal to the Hrp48 binding site (Fig. 5A). Replacing Hrp48 with Hrp48-ASO rescued stable Sxl dwell times in the presence of Unr (Fig. 5B). To test whether longer-lived Sxl-bound dwell times occur when Hrp48 is simultaneously bound to the same RNA molecule, we performed multi-color smFRET assays to simultaneously monitor binding of Hrp48-Cy5 and Sxl-dRBD5-Cy5.5 (Fig. 5C,D). These experiments confirm that Sxl-bound dwell times in physical presence of Hrp48 are longer than in its absence (Fig. 5E).

To functionally validate our findings, we performed bulk translation assays of an *msl-2* reporter mRNA that either lacks or contains the RNA sequence that competes with the Sxl and Unr binding sites (Fig. 5F). We observed that Sxl-dependent translation repression of *msl-2* is increased in a reporter mRNA that lacks the competing nucleotides. Overall, our data agrees with a model in which Hrp48 increases the stability of the Sxl·Unr·*msl-2* complex not by contributing with additional protein-protein interactions but rather by preventing inhibitory RNA structures that otherwise would compete with the Sxl and Unr binding sites and push the two proteins away from the *msl-2* 3’ UTR. Therefore, we propose that Hrp48 exerts its function in *msl-2* mRNA translation repression by acting as an ATP-independent RNA chaperone. Overall, our data provides a first molecular and biophysical explanation for observations where mutating the Hrp48 binding site decreases Sxl-mediated translation repression activity (Szostak et al., 2018).

## Discussion

Using smFRET, NMR and translation assays, we discovered multiple synergistic assembly mechanisms of a translation-repression-competent mRNP complex formed on the 3’ UTR of the *msl-2* mRNA. These mechanisms include: First, RNA-coupled folding of intrinsically disordered residues C-terminal to Sxl’s RBDs into an α-helix extends Sxl’s bound dwell time at its target site, correlating with enhanced translation repression (Fig. 2). Second, the *msl-2*_EaFb_-tandem repeat stabilizes Sxl binding either by allowing sliding between E and F sites or by permitting simultaneous binding of two Sxl proteins (Fig. 4). Third, initial Sxl binding accelerates Unr recruitment by more than two orders of magnitude (Fig. 3). Fourth, Hrp48 facilitates *msl-2*_EaFb_ site accessibility by resolving inhibitory RNA secondary structures, further prolonging Sxl bound dwell times (Fig. 5). We demonstrate that these kinetic effects are directly related to translation repression activity using reporter gene assays (Fig. 2, Fig. 5, Fig. S3). Together, our data support a model (Fig. 6) in which multiple proteins dynamically cooperate with each other and the RNA to ensure efficient *msl-2* mRNA translation repression.

**Fig. 6.**
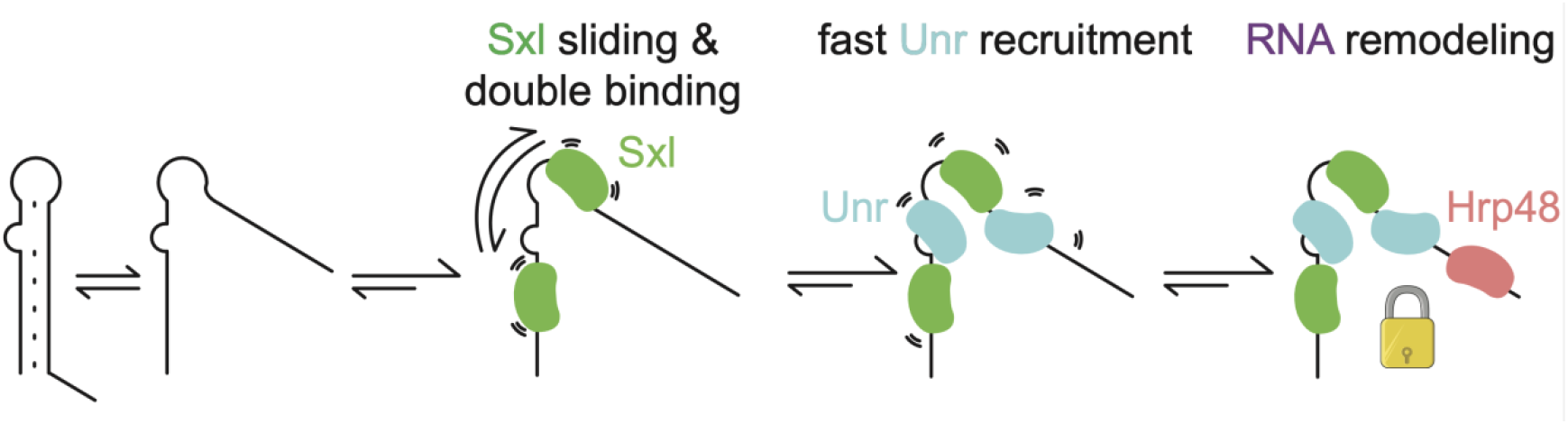
Kinetic model of *msl-2* mRNP complex formation. Different mechanisms such as sliding and double binding of Sxl to repeats of U-tracts, ultrafast Sxl-mediated Unr recruitment and RNA chaperoning by Hrp48 synergistically increase mRNP repression complex occupancy on the 3’ UTR and thus, translation repression activity.

We show that Unr binding is strongly accelerated by RNA-bound Sxl and that Sxl/Unr assembly on the *msl-2* 3’ UTR occurs hierarchically, with Sxl recruiting Unr. While a previous study revealed that Sxl and Unr cooperate via RNA-dependent protein-protein contacts that wrap around the sandwiched RNA to stabilize the mRNP complex (Hennig et al., 2014), it was unclear from a structural snapshot alone that this cooperativity also enhances Unr’s binding rate. Using smFRET, we could measure in real-time that cooperative Sxl/Unr binding is driven by increased on-rates when the other protein is already bound to the RNA (Fig. 3). Increased RNA structure- and sequence specificity can be achieved by multidomain RBPs whose RBDs act cooperatively (e.g. Duszczyk et al., 2022; Hennig and Sattler, 2015; Hollmann et al., 2020; Mackereth et al., 2011). However, how intermolecular RBDs enhance specificity through RNA-mediated cooperativity remains poorly characterized, despite its relevance in RNA biology. We propose that the ultrafast recruitment of secondary RBPs via RNA-dependent protein-protein cooperativity is a common feature of multi-RBP systems and a general mechanism in assembling transient regulatory mRNP complexes. Similar hierarchal assembly is seen in molecular machines, for example, during pre-mRNA splicing, where 15.5K binding to the U4 5’ stem loop is required to recruit Prp31, which cannot bind to 15.5K or U4 snRNA in isolation (Liu et al., 2007).

Several studies have shown that regions outside of conserved RBD boundaries, often overlooked in structural analyses, can significantly enhance RNA binding. For instance, RNA binding induces an α-helix in a disordered region C-terminal to RBM20’s RRM domain (Upadhyay and Mackereth, 2020), a splicing regulator associated with dilated cardiomyopathy (Li et al., 2010; Van Den Hoogenhof et al., 2018). Similarly, seven residues C-terminal to Sxl’s RRM2 domain are essential for translation repression (Grskovic et al., 2003). Using smFRET, NMR and ITC we clarified the molecular mechanism of this C-terminal RRM extension: while ITC and NMR revealed increased RNA affinity and RNA induced α-helix formation, smFRET showed that this affinity enhancement mainly stems from prolonged RNA-bound dwell times, which results in enhanced translation repression activity.

Although Sxl has long been known to bind adjacent E and F sites within the 3’ UTR (Bashaw and Baker, 1997; Gebauer, 1999; Kelley et al., 1997), how assembly at these poly(U) repeats is initiated remained unclear. A previous study (Hennig et al., 2014) showed that Sxl and Unr can assemble onto the *msl-2*_EaFb_ mRNA in a 2:2:1 ratio at high µM concentrations, but the necessity for both sites and its assembly mechanism remained unclear. We find that Sxl moves between the E and F-sites, increasing the retention time on the *msl-2* mRNA before dissociation and thereby enhancing the likelihood of mRNP complex assembly. While sliding has been observed for other RNA targeting complexes such as miRNA-bound Ago (Chandradoss et al., 2015; Globyte et al., 2019) and double-stranded RNA binding domains (Koh et al., 2013; Masliah et al., 2018; Tants et al., 2017), we observed with smFRET, that RBPs targeting ssRNA use sliding along the mRNA to achieve their function as well. Our data highlights that Sxl utilizes lateral movement as a key mechanism for efficient translation repression. Notably, clustering of Sxl binding sites is conserved across different *Drosophila* species (Hennig et al., 2014), with species like *D. mojavensis, D. virilis* and *D. grimshawi* even having acquired a third Sxl/Unr binding motif between sites E and F. We propose that repeats of *cis*-regulatory binding elements on UTRs represent a general evolutionary strategy to ensure robust mRNP assembly and functionality. Supporting this, a human Sxl orthologue HuR more efficiently localizes nascent CD47 to the plasma membrane via binding to the *cd47* 3’ UTR when multiple AU-rich elements, the cognate HuR binding motif, are present (Berkovits and Mayr, 2015).

Sxl and Unr are essential but insufficient for full translation repression of the *msl-2* mRNA. A third factor, Hrp48, was previously identified as a crucial component of this mRNP complex (Szostak et al., 2018), but its role in mRNP complex assembly remained unclear. Given the proximity of the Hrp48 binding motif to the E and F sites (Lomoschitz et al., 2025), we initially hypothesized that Hrp48 stabilizes the complex through additional protein-protein contacts. However, our data reveal that Hrp48 functions instead as a ‘doorstop’, preventing RNA hairpin formation and thereby enhancing retention of Sxl and Unr to their target site. This suggests that Hrp48 can chaperone RNA structure through dynamic, reiterative binding, stabilizing the mRNP complex. This mechanism resembles ribosome assembly, where secondary binding proteins increase the stable association of primary binding proteins (Duss et al., 2019). We further show that RNA-bound dwell times of Sxl in presence of Unr and Hrp48 can reach up to 30 s under physiological conditions (Fig. 5B). Since translation initiation itself takes ∼ 30 s on average (Koch et al., 2020; Shah et al., 2013), such a stable repressive mRNP complex likely renders *msl-2* translation effectively impossible. To our knowledge, this is the first evidence that a ubiquitous RBP like Hrp48 can act as an ATP-independent RNA remodeler to promote mRNP complex assembly.

In conclusion, we propose a model in which diverse and likely widespread mechanisms – including RBP sliding along RNA, rapid recruitment of secondary proteins and RNA remodeling – cooperate to achieve sufficient mRNP complex stability for proper function. To our knowledge, this is the first comprehensive study to dissect the dynamic assembly pathway of a functional, transient multiprotein-RNA complex, moving beyond the classical view of single RBP-RNA interactions or stable molecular machines. Overall, we provide a framework and toolbox to study dynamic, multicomponent protein-RNA complex assembly in real-time, showcasing how powerful and informative this approach can be for unravelling the mechanisms of RNP formation and function.

While this study uncovers multiple mechanisms of mRNP assembly that enhance translation repression, it does not directly address how translation of the *msl-2* mRNA is inhibited. It has been proposed that the Sxl·Unr·Hrp48·mRNA complex blocks translation by interacting with components of the 43S preinitiation complex, preventing access to the 5’ region and therefore start codon scanning (Gebauer et al., 2003), with a potential interaction between eIF3d and Hrp48 suggested (Szostak et al., 2018). However, alternative mechanisms, such as sequestration of the *msl-2* mRNA into biomolecular condensates inaccessible to ribosomes, could be a viable mechanism to prevent 43S PIC recruitment. Notably, Hrp48 is known to readily undergo liquid-liquid phase separation in other contexts (Bose et al., 2022). Addressing these possibilities would require either reconstituting this system including the 43S complex for cryo-EM studies or investigating condensate formation and its link to translation repression – both beyond the scope of this study.

## METHODS

### Plasmids

Single cysteine mutants of Sxl-dRBD3 (C154S and S207C) were derived from pET-24d(+) (Hennig et al., 2014) using site-directed mutagenesis (Braman et al., 1996) and Q5 DNA polymerase (NEB). Sxl-dRBD5 (122-305) was derived from full-length Sxl and inserted via overlap extension PCR (Bryksin and Matsumura, 2010) into a pET-24d(+) vector. Single cysteine mutations of Sxl-dRBD5 (C154S/S207C or N-terminal Cys/ C154S) were introduced with site-directed mutagenesis. For AviTag-Sxl-dRBD5 (N-terminal Cys/C154S), the Avi-tag peptide sequence (GLNDIFEAQKIEWHE) (Beckett et al., 1999) with a (G_3_SG_3_SG_2_S) linker was fused N-terminally. Sxl constructs comprise an N-terminal His_6_-tagged Trx with a TEV cleavage site. Unr full-length (CG7015-PA) with a carboxy-terminal His_6_ was cloned from a SL2 cDNA library using overlap extension PCR into a pETM11 (derived from pBR322; G. Stier) as described (Hollmann et al., 2023). A single cysteine mutant of Unr-CSD1 including a C-terminal Tryptophan to aid spectroscopic protein quantification (186-252; C23S and W253) was derived using site-directed mutagenesis (Hennig et al., 2014). Hrp48 constructs as described in Bose et al., 2022 were modified to contain a TEV cleavage site C-terminal of the eGFP tag to allow StrepTagII removal, inserted via primer extension PCR. For Hrp48 constructs, the ybbR peptide tag (DSLEFIASKLA) (Yin et al., 2005) was introduced N-terminal to the Hrp48 CDS with overlap extension PCR, followed by 5′-end phosphorylation with T4 PNK (NEB) and blunt ligation with T4 DNA ligase (NEB). Translation reporter constructs of *msl-2* were constructed via NEBuilder® HiFi assembly (NEB). Constructs contain a T3 promoter with a 5’ UTR from the rLuc plasmid, an open reading frame that consists of N-terminal 5 x alfa-tags (SRLEEELRRRLTE) (Götzke et al., 2019) separated by a linker (P(G_4_S)_3_P) fused to a firefly luciferase gene, and the minimal 3’ UTR regions from *msl-2* (CG3241-RA) with an extension (3702-3761). A deletion construct of the 3’ UTR extension (3750-3761) was derived via HiFi assembly using DNA primers that encode the minimal 3’ UTR region. Cloned plasmids were transformed into Top10 via electroporation.

### Protein expression and purification

Sxl (dRBD3, dRBD5) and Unr proteins (CSD1) were expressed in *E. coli* BL21 (DE3) in terrific broth (TB) with 1 mM IPTG at OD600 = 1.0 at 20 °C overnight. Unr full-length was expressed and purified as described (Hollmann et al., 2023). Harvested cells were resuspended in lysis buffer A (50 mM Hepes/NaOH, 500 mM NaCl, 1M urea, 2 mM MgCl_2_, pH 7.5) supplemented with cOmplete protease inhibitor (Roche), benzonase and lysed by passing through a microfluidizer 4 times. The cleared lysate was loaded onto a 5 ml HisTrap FF (Cytiva) and eluted with a gradient (lysis buffer A + 500 mM Imidazole) after 7 column volumes (CVs) of high-salt wash (50mM Hepes/NaOH, 1 M NaCl, 2 M LiCl, pH 7.5) and 7 CVs of wash with lysis buffer A. Sxl and Unr proteins were cleaved overnight with in-house His_6_-TEV protease at 4 °C in 2 l of dialysis buffer (50 mM Hepes/NaOH, 50 mM NaCl, pH 7.5) using dialysis tubings with appropriate cut-offs. After cleavage, proteins were loaded onto a HisTrap, washed with at least 5 CVs of dialysis buffer and collected as either flow-through, or unspecifically bound proteins during isocratic step elutions. Sxl proteins (dRBD3, dRBD5) and Unr-CSD1 were buffer exchanged using a HiPrep 26/10 desalting column to heparin buffer A (50 mM Hepes/NaOH, 50 mM NaCl, pH 7.5) and heparin buffer 3A (20 mM NaPi, 50 mM NaCl, pH 6.0), respectively. Sxl proteins (dRBD3, dRBD5) and Unr-CSD1 were loaded onto a heparin HP 5 ml and eluted using either heparin buffer B (50 mM Hepes/NaOH, 1000 mM NaCl, pH 7.5) or heparin buffer 3B (50 mM NaPi, 1500 mM NaCl, pH 6.0), respectively. In a final step, all proteins were buffer exchanged against SEC buffer (50 mM Hepes/NaOH, 50 mM NaCl, pH 7.5) using a S75 16/600 (Cytiva) size exclusion column and concentrated using Amicon concentrators with appropriate molecular weight cut-offs. Protein quality was assessed via SDS-PAGE and Coomassie staining and the concentration was determined photometrically. Hrp48 was purified as described (Bose et al., 2022) with slight modifications. Hrp48 constructs (WT & Hrp48-ybbR) were transformed into DH10Bac for bacmid production. 3 ml Sf-9 cells (0.3·10^6^ cells/ml) were transfected with 10 µg bacmids using FuGene HD transfection reagent (Promega) and the first virus generation (V0) was harvested 72 h post-infection (PI), checking for protein expression under a fluorescence microscope. The second virus generation (V1) was obtained by harvesting 72 h PI with V0 (3 ml into 25 ml Sf-9 at 0.5·10^6^ cells/ml). For large-scale expression, Sf-21 cells (0.7-1.0·10^6^ cells/ml) were infected with V1 (1:100) in Sf-900 III SFM medium (Thermo Fisher) and Hrp48-expressing cells were harvested 72 h PI. Cells were flash-frozen in liquid nitrogen and stored at -20°C. Thawed cells were resuspended in binding buffer (50 mM Tris/HCl pH 7.5, 500 mM NaCl, 1 mM EDTA) supplemented with 0.01 % Triton X-100, 1 x cOmplete protease inhibitor, benzonase and 3 mM MgCl_2_ and lysed by passing it through a microfluidizer 3 times. Hrp48 constructs were loaded onto a Strep-Tactin column (5 ml, IBA Lifesciences), washed with 5 CVs binding buffer and subsequently eluted with elution buffer (binding buffer + 2.5 mM desthiobiotin). Pooled fractions were cleaved overnight at 4 ºC with His-tagged TEV protease in dialysis buffer (50 mM Tris/HCl pH 7.5, 500 mM NaCl) and subsequently loaded on a Strep-Tactin column. The flow-through was then gel filtrated on a Superdex 10/300 GL increase S200 into Hrp48 storage buffer (20 mM Tris/HCl pH 7.5, 300 mM NaCl, 2 mM MgCl_2_, 0.5 mM TCEP, 5 % (v/v) glycerol). Pooled fractions were concentrated using Amicon concentrators, aliquoted, flash frozen in liquid nitrogen and stored at -70 ºC.

For NMR samples, isotopically labelled Sxl-dRBD5 (^13^C, ^15^N, ^2^H) and Unr-CSD1 (181-252, ^2^H) were expressed in *E. co*li BL21 (DE3) according to Li and Byrd, 2022. Cells were harvested by centrifugation (15 min, 4 °C, 9000 g, JLA 8.1000, Avanti J-26 XP). Cells were resuspended in lysis buffer (50 mM NaPi, 300 mM NaCl, 1 mM DTT, pH 7.2), and lysed using a microfluidizer, followed by 1 h centrifugation at 16,500 g at 4 °C. The supernatant was loaded on a 5 ml HisTrap HP column and the protein eluted with lysis buffer containing 500 mM imidazole. TEV protease cleavage was carried out over night during dialysis against 50 mM NaPi, 300 mM NaCl, 10 mM Imidazole, 1 mM DTT, pH 7.2 at 4 °C. IMAC was repeated to remove the cleaved purification tag. Purified protein samples were stored at -80 °C.

### Protein fluorophore conjugation

Labeling of Sxl and Unr-CSD1 constructs was performed as described earlier (Duss et al., 2018). In short, sulfo-Cy3-, sulfo-Cy5- or sulfo-Cy5.5-maleimide fluorophores were conjugated to constructs bearing a single cysteine. Sxl-dRBD3 was labeled with sulfo-Cy5 and Sxl-dRBD5 with sulfo-Cy5, sulfo-Cy5.5 or sulfo-Cy3. First, 1 mg of protein was fully reduced in 500 µl of labeling buffer A (100 mM Na_2_HPO_4_/KH_2_PO_4_ pH 7.0, 100 mM NaCl) overnight and then precipitated using 70 % (w/v) ammonium sulfate and centrifugation after 20 min incubation. After two washes with ice-cold labeling buffer A (+ 70 % ammonium sulfate), the protein pellet was dissolved in 500 µl labeling buffer A1 (labeling buffer A + 6 M urea). To this, 1 mg of either sulfo-Cy5-maleimide (Cytiva), sulfo-Cy3-maleimide or sulfo-Cy5.5-maleimide (Lumiprobe) was added and incubated at room temperature for 1 h under gentle agitation and light protection. To remove excess dye, the reaction was quenched with β-mercaptoethanol (0.5 %; 72 mM) and loaded onto pre-equilibrated NAP-5 columns (Cytiva). Collected fractions were analyzed with SDS-PAGE and imaged using a Typhoon FLA 7000 and subsequent Coomassie staining. Desirable fractions were pooled and refolded via 15-fold dilution using either heparin buffer 3A (20 mM Tris/HCl pH 7.6, 20 mM NaCl, 0.5 mM EDTA) for labeling of Sxl or heparin buffer 4A (20 mM NaPi pH 6.0, 20 mM NaCl, 0.5 mM EDTA) for labeling of Unr-CSD1. Diluted pooled fractions were passed onto a 1 ml heparin FF (Cytiva) and washed with 10 CVs before eluting with heparin buffer 3B (20 mM Tris/HCl pH 7.6, 1 M NaCl, 0.5 mM EDTA) or heparin buffer 4B (20 mM NaPi pH 6.0, 1 M NaCl, 0.5 mM EDTA), respectively. Collected fractions were assessed for free dye using SDS-PAGE and fluorescent imaging.

### Sfp1-mediated ybbR CoA-fluorophore labeling

Hrp48-ybbR (10 µM) was natively labeled using 7.5 µM Sfp1-mediated coupling of 3-fold molar excess of CoA-Cy5 (30 µM) in Hrp48 storage buffer supplemented with 10 mM MgCl_2_. The reaction was incubated for 30 min at room temperature, followed by overnight incubation at 4 ºC under slight agitation using a rotating wheel. The labeled protein was purified by passing it through a Zeba™ spin desalting plate (7K MWCO).

### *In vitro* biotinylation

Cy3-labeled AviTag-Sxl-dRBD5 was dialyzed against reaction buffer (10 mM Tris/HCl, 50 mM potassium (L)-glutamate, pH 7.5) and biotinylated *in vitro* using 10 µM of protein that was conjugated to d-biotin (in DMSO) at 15-fold molar excess using 7 µM BirA ligase in 1 x birA reaction buffer (50 mM Tris/HCl, 28 mM potassium (L)-glutamate, 5 mM MgCl2, 2 mM ATP). The reaction was incubated at room temperature for 2 h under slight agitation. Unreacted biotin was removed from the reaction using a heparin HP 1 ml column. Loaded protein was washed with 10 CVs of 1 x birA reaction buffer, 10 CVs heparin 3A and eluted with heparin 3B. Successful biotinylation was validated with a Tamavidin2 SDS-PAGE shift assay.

### Protein concentration and labeling efficiency

Protein concentration of labeled proteins was determined via Bradford and cross-validated photometrically. Maximum absorbance values of the individual dyes conjugated to the proteins was also determined spectroscopically. Labeling efficiency was then calculated as the ratio of fluorophore concentration and protein concentration.

### *In vitro* transcription and capping

*In vitro* transcriptions were carried out using pre-annealed DNA oligonucleotides (for lengths up to 120 bp) as DNA templates for run-off transcription. DNA strands (45 µM each) were annealed at 95 °C for 5 minutes and cooled down to room-temperature at 0.1 °C/s in 1 x DNA annealing buffer (10 mM Tris/HCl pH 7.5, 50 mM NaCl, 1 mM EDTA). Larger DNA templates (<1000 bp) for *in vitro* transcription were generated using the PCR product of overlapping primer fragments (Tian and Das, 2017) giving rise to the full construct. DNA templates consisted of a T7 promoter followed by one or two guanines, an F2 sequence (5’-AACCACUCCAAUUACAUACACC-3’) for labeling the RNA, and the RNA sequence of interest. The AC-rich sequence (5’-ACUACCACCACCCAACCAACACACC-3’) for subsequent immobilization was placed at the 5’ or 3’ end (see Table S1). For Hrp48 binding studies, RNA was labeled via the AC-rich sequence and immobilized via the F2 sequence.

*In vitro* transcription reactions were performed at 37 °C using 5 mM DTT, 0.25 mg/ml in-house T7 RNA polymerase, 0.02 U/µl thermostable inorganic pyrophosphatase (NEB), and 0.1 µM DNA template in 1 x transcription buffer (40 mM Tris/HCl pH 7.9, 1 mM spermidine, 0.01 % Triton X-100). For each *in vitro* transcription, MgCl_2_ and NTP concentrations were optimized ranging from 20-80 mM and 20-60 mM (total rNTP), respectively. Preparative-scale reactions were quenched after 4 h with RNA loading dye (95 % formamide, 0.01 % bromophenol blue, 0.01 % xylene cyanol FF, 0.25 mM EDTA) and the product was separated from impurities via denaturing PAGE (6 M urea) and purified using the crush and soak method (Petrov et al., 2013), followed by ethanol precipitation and resuspension in nuclease free water. Transcription templates that were generated from linearized plasmids (*msl-2* reporter constructs with EcoRI-HF (NEB); rLuc reporter with SacI-HF(NEB)) were purified with phenol/chloroform/isoamyl alcohol extraction and ethanol precipitation. Long-template transcription reactions from linearized plasmids were transcribed with a T3 RNAP (NEB) at 37 ºC for 2 h in 1 x transcription buffer and post-transcriptionally capped using the Vaccinia Capping System (NEB) according to manufacturer’s instructions.

### *Drosophila* embryo extract preparation

*Drosophila* embryo extracts were prepared as described (Gebauer, 1999) with slight modifications. 90-110 min synchronized *D. melanogaster* embryos were collected by exchanging yeasted apple agar plates 3 times every hour prior to final collection. Embryos were washed extensively with EW buffer (120 mM NaCl, 0.04 % Triton X-100) using stacked sieves, dechorionated for 3 min in a 1:1 mix of water and sodium hypochlorite solution (6-14 %). Embryos were again washed extensively with EW buffer and subsequently washed with DE buffer (10 mM HEPES pH 7.4, 5 mM DTT). Packed embryos were diluted with an equal volume of DEI buffer (DE buffer + 1 x cOmplete protease inhibitor) and lysed with 20 strokes in a dounce homogenizer. The crude extract was centrifuged at 13,300 g for 10 min and cytoplasmic extract was obtained as the interphase between pelleted nuclei/cell debris and lipid layer. Cytoplasmic extract was aliquoted, flash frozen in liquid nitrogen and stored at -70 ºC. Total protein amount was measured via Bradford using BSA as a standard.

### Translation assays

*In vitro* translation assays were adapted from (Fukaya and Tomari, 2011; Gebauer, 1999). In a 10 µl reaction, 20 nM of capped renilla luciferase mRNA, 20 nM capped firefly luciferase-containing mRNA, 4 mg/ml *Drosophila* embryo extract, 3 µl of 40 x mix, 50 mM potassium acetate and 2 mM Mg(OAc)_2_ were incubated for 90 min at 25 ºC. Sxl was added to the reaction at a final concentration of 1.2 µM. Firefly and renilla luciferase translation levels were obtained on a Biotek Synergy H4 plate reader from 4 µl of translation reaction using the Dual-Luciferase® Reporter Assay System (Promega) according to manufacturer’s instructions. Different conditions were measured in triplicates and normalized translation of condition *i* was obtained as following.

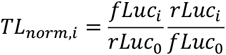

### NMR spectroscopy

Purified proteins were dialysed against NMR buffer (10 mM KPi, 50 mM NaCl, 1 mM DTT, pH 6.0) and subsequently concentrated to 300 µM final concentration. The ^15^N,^13^C,^2^H-Sxl-dRBD5: ^2^H-Unr-CSD1: ^1^H-*msl-2*_Fb_-RNA complex was assembled stepwise. To trace binding saturation, TROSY experiments were carried out after addition of each component of the complex. *Msl-2*_Fb_ RNA was added in 1.2-fold excess with respect to Sxl-dRBD5 and Unr-CSD1 was added in 1.5-fold excess. After final assembly of the complex, the diluted sample was concentrated again to 300 µM to acquire triple resonance experiments.

All NMR spectra were recorded on Bruker Avance IIIHD 700 MHz, 900 MHz, and 1000 MHz NMR spectrometers, equipped with cryogenically-cooled triple-resonance probes at a temperature of 298 K.

For protein resonance assignments, BEST-TROSY-based double and triple resonance through-bond experiments were recorded (Favier and Brutscher, 2011; Lescop et al., 2007). Apodization weighted sampling was used for acquisition of triple resonance spectra (Simon and Köstler, 2019). NMR data were processed using NMRPipe (Delaglio et al., 1995) or in-house routines and analyzed with Cara (https://cara.nmr.ch), NMRViewJ (Johnson and Blevins, 1994) and TALOS+ (Shen et al., 2009). Secondary chemical shifts were derived according to Wishart and Sykes, 1994. Chemical shift perturbations (CSPs) were calculated according to 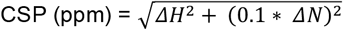 (Williamson, 2013).

### RNA 3’ labeling

The *msl-2*_Fb_ RNA (see Table S1) used for fluorescence polarization assays was labeled at the 3’ end with Fluorescein-5-thiosemicarbazide (FTSC, Cayman chemical company)(Qiu et al., 2015). RNA (6.3 µM) was oxidized with 10-fold molar excess of sodium periodate (Sigma-Aldrich) in labeling buffer (63 mM sodium acetate pH 5.6) for 90 min at 25 °C in the dark. The reaction was quenched with 120 µM of sodium sulfite (Sigma-Aldrich) for 15 min at 25 °C. Oxidized RNA was labeled with 167 µM FTSC for 3 h at 37 °C. Labeled RNA was ethanol precipitated and washed 3 times with 70 % ethanol before resuspending the dried pellet in water. Labeling efficiency (74 %) was determined photometrically in 0.1 M NaOH.

### Fluorescence polarization assay

3.5 nM of FTSC labeled RNA was incubated with different Sxl concentrations in FP buffer (Tris/HCl pH 7.5 10 mM, KCl 20 mM, NaCl 25 mM, NP-40 0.005 % (v/v)) for 20 min prior to measurement. 30 µl of each condition was measured in triplicates in a 384-well black opaque OptiPlate (Fisher Scientific) on a Biotek Synergy H4 plate reader with excitation and emission of 485 and 528 nm, respectively. Affinities were obtained from fit curves in python to normalized fluorescence polarization data as described (Jagtap et al., 2023), where X denotes the logarithmic protein concentration in nM:

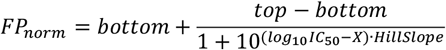

### Isothermal titration calorimetry

ITC measurements were performed using a NanoITC LV (TA Instruments) calorimeter at 298 K. Prior experiments, Sxl-dRBD3, Sxl-dRBD5 constructs and U10-mer RNA were dialyzed against different buffers (50 mM Tris/HCl pH 7.5, 50 or 150 mM KCl). Protein samples were adjusted to 60 µM and U10-mer RNA was set to a final concentration of 600 µM. Samples were degassed using a degassing station (TA instruments). The protein sample was injected into the sample cell (300 µl) and 50 µl U10-mer were loaded into the injection needle. Multiple injection experiments were carried out, each consisting of 40 injections of 1 µl titrant with 200 s spacing between each injection and a stirring rate of 300 rpm. Samples were measured in duplicates. For data analysis, the software NanoAnalyze (TA Instruments) was used.

### Mass photometry

Mass photometry experiments were performed on a Refeyn_MP_. While Hrp48 and Unr were prepared as 5 µM stocks, Sxl-dRBD5 was prepared as a 25 µM stock in 1 x e55 (10 mM Tris/HCl pH 7.5, 20 mM KCl) buffer. Proteins were incubated for 5 min at room temperature by mixing them in a 1:1 ratio. Protein mixtures were then diluted 6.3-fold in 1 x e55 buffer and 1 µl was added to 19 µl of degassed and filtered PBS (137 mM NaCl, 2.7 mM KCl, 10 mM Na_2_HPO_4_, and 1.8 mM KH_2_PO_4_) buffer residing on the sample carrier slide and mixed by pipetting. Particle mass was detected as the interference of scattered light with the surface-reflected light. Mass calibration was performed by using BSA and IgG.

### Single-molecule fluorescence microscopy

Passivation of glass slides with NHS-ester mPEG/biotin-PEG (Laysan Bio Inc.) coating was performed as described (Chandradoss et al., 2014). Passivated glass slides were stored under N_2_ gas at -20 °C and right before experiments, a second round of PEGylation with MS4-PEG (Thermo Fisher) was performed. For ALEX experiments, glass slide passivation with PEG was performed at cloud-point conditions as described (Zhang et al., 2016). The second round of PEG coating was done with NHS-acetate.

Single-molecule imaging experiments were conducted using a custom-built objective based (CFI SR HP Apochromat TIRF 100×C Oil) TIRF microscope (by Cairn Research: https://www.cairn-research.co.uk/) with an iLas targeted laser illuminator for uniform widefield illumination. For a 2-color set-up (Cy3 & Cy5), FRET donor dyes were excited using a diode-based (OBIS) 532 nm laser from a Cairn Multiline laserbank and FRET acceptor dyes were directly excited using a diode based (Omicron LuxX) 638 nm laser. Donor signal was separated from acceptor fluorescence using a dichromatic beam splitter (T635LPXR, Chroma). Emission light of Cy3 was filtered using a bandpass excitation filter (ET585/65m, Chroma) and Cy5 emission was attenuated using a bandpass emission filter (ET700/75m, Chroma). For 3-color set-ups (Cy3, Cy5 & Cy5.5), a dichroic beam splitter separated Cy3 and Cy5/Cy5.5 emission (T635LPXR) and a filter cube (ET667/30m, T685LPXR, ET720/60m, Chroma) was used to separate Cy5 from Cy5.5 emission signals. Images were acquired on Prime95B sCMOS cameras (Teledyne Photometrics).

For single-molecule experiments, *in vitro* transcribed RNA was annealed with a Cy3-DNA for labeling and a DNA duplex for immobilization. The duplex consists of a biotinylated-DNA and an oligonucleotide strand that connects RNA with the biotin-DNA. The annealing was performed for 15 minutes at 60 °C in 1 x e55 buffer, followed by a slow cool-down (0.1 °C/s) in a thermal cycler.

Immobilization of individual RNA molecules was achieved by incubation of the flow chamber with NeutrAvidin for 5 minutes, followed by extensive washing with 1 x e55, and incubation with RNA for 10 min (80-200 pM). After a high-salt wash (50 mM Tris/HCl pH 7.5, 150 mM KCl, 20 mM NaCl, 14 mM MgCl_2_), a combination of labeled and unlabeled proteins in 1 x e55 buffer (+ NaCl to desired ionic strengths) with TSY (2 mM) as a triplet state quencher, PCA (2.5 mM) and PCD (0.42 U/µl) as an oxygen scavenging agent was injected into the flow chamber and imaged (Duss et al., 2019, 2018).

Acquisitions were performed at a laser power density of 0.86 kW·cm^-2^ (100 % 532 nm laser power density; on the basis of output power) at an evanescent field depth of 250 nm with 100 ms exposure time and at a temperature of 21 °C if not stated otherwise.

For 2-color experiments at 21 °C, binding of Cy5 labeled Sxl constructs was measured at an ionic strength of 45 mM (adjusted with NaCl) in 1 x e55 buffer. If not stated otherwise, used concentrations were: Sxl-dRBD3 (40 nM total, 84 % labeling efficiency), Sxl-dRBD5 (batch 1: 40 nM total, 70 % labeling efficiency or batch 2: 34.5 nM total, 82 % labeling efficiency). Binding of Unr-CSD1-Cy5 was monitored at 19 mM ionic strength at 40 nM total protein concentration (65 % labeling efficiency). Movies with > 10 min length were recorded with a laser power of 80 % (for Sxl-dRBD3/5-Cy5), and 70 % (for Unr-CSD1) 532 nm. Unlabeled Unr-CSD1 or Sxl-dRBD3/5 was added at a final concentration of 1 µM. Binding of Sxl-dRBD5-Cy5 or Hrp48-Cy5 to *msl-2*_EaFbH48_ or *msl-2*_3’UTR_ was measured in 1 x e55 at 150 mM ionic strength (adjusted with NaCl) and 29 °C at 100 % 532 nm laser power density if not stated otherwise. For 2-color experiments at 29 °C, unlabeled proteins were added at 400 nM concentrations.

In all 3-color experiments, if not stated differently, 100 % 532 nm laser power density was used. Binding of Sxl-dRBD5-Cy5 (40 nM total) and Unr-CSD1-Cy5.5 (200 nM total, 71 % labeling efficiency) was recorded at 200 ms exposure time. Sxl-dRBD5-Cy5.5 (40 nM total, 99 % labeling efficiency) and CSD1-Cy5 (200 nM total) was measured at 100 % 532 nm laser power density and a 200 ms exposure time. 3-color experiments with Hrp48-Cy5 (95 nM total, 65 % labeling efficiency) and Sxl-dRBD5-Cy5.5 (30 nM total) were recorded at 29 °C with 150 mM ionic strength at 100 % 532 nm laser power density and a 100 ms exposure time.

For ALEX experiments, 100 ms exposure time for each laser was used with a laser power density of 80 % for 532 nm and 15 % for 638 nm (0.17 kW·cm^-2^), respectively. 40 µg/ml BSA were supplemented to avoid stickiness. Sxl-dRBD5-Cy5 was added at 40 nM total protein concentration.

In 1D diffusion assays, biotinylated AviTag-Sxl-dRBD5-Cy3 was immobilized with 100 pM labeled protein and 60 nM Cy5-labeled *msl-2*_EaFb_ binding was monitored using 60 % 532 nm laser power density at an exposure time of 100 ms. For Cy5 labeling of *msl-2*_EaFb_, the RNA was annealed to a Cy5-labeled DNA oligonucleotide complementary to the F2 site and a second DNA oligonucleotide complementary to the AC-rich overhang for 15 min at 60 ºC before slow cooling (0.1 °C/s) to room temperature.

### Single-molecule data processing and event assignment

Movies were recorded as tiff stacks and preprocessed using MetaMorph (Molecular Devices). Stacks were separated into individual channels and assembled into tiles suitable for further processing in Spartan (Juette et al., 2016). After tile generation, movies were drift corrected if necessary, using cross-correlation of the donor signal intensity using Python scripts. In Spartan, channels of drift corrected tiles were aligned with an acquired beadslide before exporting the traces. Exported traces were analyzed further with MATLAB (Lapointe et al., 2022; Qureshi and Duss, 2025) and picked traces in Spartan were manually curated and inspected for absolute fluorescence intensity values or single photobleaching steps to ensure single-molecule time traces. Binding events in picked traces were then assigned via trace specific thresholding of transfer efficiencies ⟨*E*⟩ = *I*_*A*_/(*I*_*A*_ + *I*_*D*_), where *I*_*A*_ denotes the fluorescence intensity of the FRET acceptor and *I*_*D*_ the intensity of the FRET donor. Due to spectral bleed-through of the Cy5 into Cy5.5 channel in 3-color measurements, binding events of Cy5.5 and Cy5 were assigned via trace-specific thresholding of the Cy3 to Cy5.5 FRET (Qureshi and Duss, 2025). In 2-color measurements with multiple FRET states, high and low FRET states were assigned via trace-specific thresholding based on transfer efficiencies.

### Quantification and statistical analysis

#### Co-binding events

In 3-color (Fig. 3, 5) or ALEX (Fig. 4) datasets, co-binding events were defined as temporally overlapping binding events (e.g. Cy5 and Cy5.5 binding events), where rebinding events would also count as co-binding events. A co-binding event in which Sxl binds first (Cy5) and Unr second (Cy5.5) would be registered as “Sxl first”. The relative ratio of “Sxl first” would be computed based on all co-binding and re-binding events measured. The arrival time of a second factor in a co-binding event would then be evaluated based on the time delay between the first factor binding and the second factor binding. In re-binding of the second factor events the time delay in between would be computed as another arrival time of the second factor. Binding events per unit of time are then computed by calculating the inverse of the arrival time.

#### Generation of 1D histograms and transition density plots

Both donor and acceptor traces were background corrected in Spartan after a single FRET donor photobleaching occurred. Traces were then further processed and corrected for donor emission bleed-through (α) and for different quantum yields of donor and acceptor dyes (γ factor). After processing, traces were exported in Spartan and 1D FRET histograms were computed from Spartan traces in tMaven (Verma et al., 2024). Fits of gaussian curves were performed using Python scripts by fitting a linear combination of up to two gaussians to the probability distribution of the transfer efficiencies derived from tMaven (unfiltered traces). Transition density plots were also computed from tMaven using default settings.

#### Maximum dwell time per trace versus all dwell times

In order to quantify and compare the effect of various combinations of other unlabeled proteins on the bound dwell times of a labeled protein, we plotted the longest dwell time per trace instead of plotting all the dwell times per trace. In absence of unlabeled proteins, each observed bound dwell of the labeled protein corresponds to only this protein bound to the RNA and thus, all observed dwells were used in data evaluation. However, in presence of unlabeled proteins, the observation of a binding event only reports on the presence of the labeled protein bound to the RNA but does not report on whether also other unlabeled proteins are co-bound. In order to enrich in protein binding events of the labeled protein at which another unlabeled protein is co-bound, we therefore plotted only the longest dwell time per trace(Duss et al., 2019, 2018).

#### On/Off-rate calculation

To determine on- and off-rates (*k*_*i*_), the cumulative distributions of all unbound and bound dwell times respectively were fit to a linear combination of up to two decaying exponentials, in which the sum of coefficients (i.e. populations of each exponential; *a*_*i*_) needs to sum up to 1. Optimal parameters were derived by non-linear least squares fits.

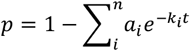

#### Figure generation

Structures were visualized with ChimeraX-1.8 and all figures were prepared using MATLAB R2019a, Excel, Python and Adobe Illustrator.

### Data and code availability

The chemical shift assignments of Sxl-dRBD5 in the free state, bound to U10-mer RNA and in complex with Unr-CSD1 and the *msl-2*_Fb_ RNA have been deposited to the Biological Magnetic Resonance Bank (BMRB) with the accession codes 52947, 52948, and 52952, respectively. The single-molecule data files are available upon request. The software used for single-molecule data processing and analysis is published (Juette et al., 2016; Verma et al., 2024) and freely available online. Additional scripts for downstream processing were made available publicly previously under an open-source license (Lapointe et al., 2022; Qureshi and Duss, 2025).

## Supporting information

Supplementary_Material_Payr_et_al_2025

## ACKNOWLEDGMENTS

M.P. would like to thank Alessandro Dulja for help in preparing *Drosophila* embryo cell-extract, Mainak Bose for sharing initial Hrp48 aliquots and Pravin Jagtap for providing full-length Unr aliquots, Cairn Research and Kavan Gor for the design and set-up of the microscope, Karine Lapouge for assistance in mass photometry experiments, Nusrat Qureshi for preparing reagents for single molecule imaging, and the Duss group for useful scientific discussions. We also thank Tanit Guitart and Fátima Gebauer for helpful discussions and providing us with plasmids and protocols. M.P. acknowledges support by a Boehringer Ingelheim Fonds PhD Fellowship. O.D acknowledges support from the FEBS Excellence Award and the European Molecular Biology Laboratory. J.H. gratefully acknowledges support by the European Molecular Biology Laboratory and the Deutsche Forschungsgemeinschaft (DFG) via grant no. 508497078.

## AUTHOR CONTRIBUTIONS

Conceptualization, M.P., J.H. and O.D..; Methodology, M.P. (smFRET, translation assays), O.D. (smFRET), J.M. (NMR, ITC), J.H. (NMR, ITC); Formal Analysis: M.P., J.M. (NMR, ITC), J.H. (NMR); Software: E.M.G.; Investigation, M.P., J.M.; Writing—original draft, M.P.; Writing—review & editing, M.P., O.D. and J.H.; Project Administration, M.P., J.H., O.D.; Funding acquisition, J.H.; Resources, J.H. and O.D.; Supervision, J.H. and O.D.

## SUPPLEMENTAL INFORMATION

**Document S1. Figures S1–S9 Table S1. Sequences**

