## Supplementary_Material_Payr_et_al_2025 for "Real-time tracking of mRNP complex assembly reveals various mechanisms that synergistically enhance translation repression"

### SUPPLEMENTAL INFORMATION

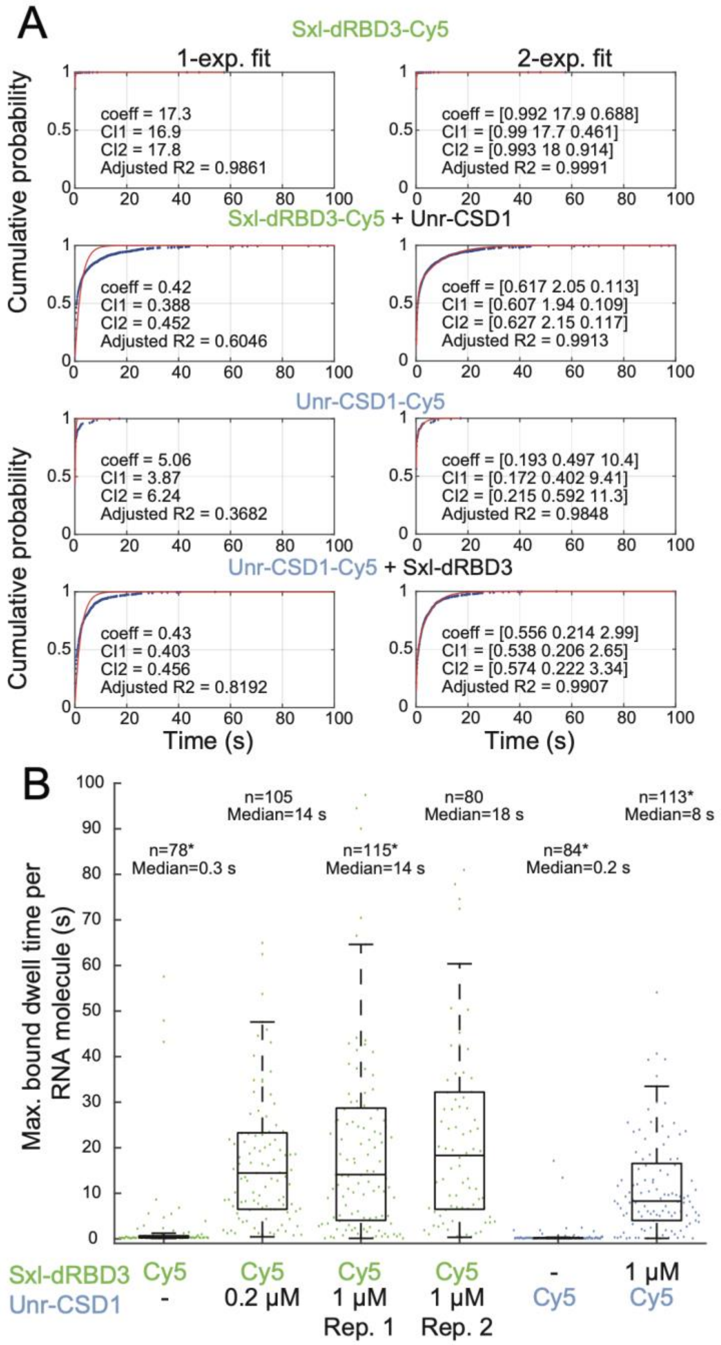


**Fig. S1. Comprehensive kinetic information of Sxl-dRBD3 and Unr-CSD1 binding to *msl-2*$_{Fb}$ mRNA, related to Fig. 1.** (A) Fits of all dwell times to a linear combination of up to two decaying exponentials of the form $p_i e^{-k_i t}$, with $k_i$ being the $i^{th}$ on- or off-rate constant and $p_i$ being the population of the respective exponential term. The "coeff" array describes the exponent for a 1-exponential fit. For a 2-exponential fit, the "coeff" array denotes the population of the first exponential, the exponent of the first exponential and the exponent of the second exponential. CI1 and CI2 describe the lower and upper 95 % confidence intervals of the fit, respectively. Fitted lifetimes for a single and double-exponential fit for dRBD3-Cy5 and CSD1-Cy5 bound dwell times are shown as scatter dots with the dot size representing the population size. (B) Replicates of dRBD3-Cy5 binding in presence of different unlabeled Unr-CSD1 concentrations show that the cooperative effect is also present at lower Unr concentrations. Replicates marked with asterisks are shown in Fig. 1.

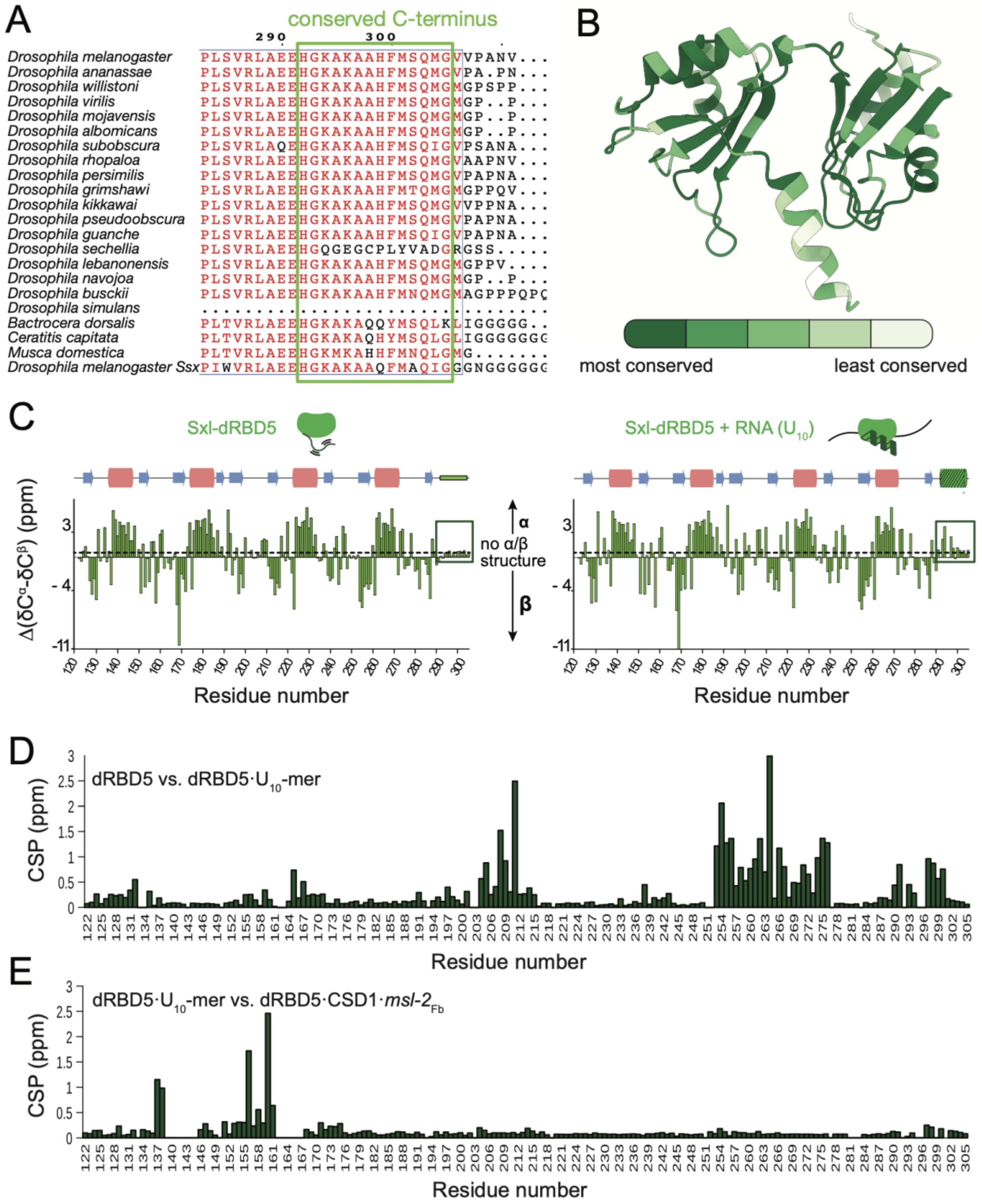


**Fig. S2. Conservation of Sxl residues involved in non-canonical RNA binding, related to Fig. 2.** (A) Multiple sequence alignment of Sxl orthologs of different species of the Drosophilidae family and other dipteran species shows a conserved C-terminal region (Sievers et al., 2011). (B) Sequence conservation of Sxl mapped onto the predicted structure of Sxl-dRBD5 (residues 122-305). (C) Secondary chemical shift perturbations of Sxl-dRBD5 in isolation (left) or in complex with $U_{10}$-mer RNA. (D) C-terminal residues of dRBD5, that are within the α-helix, show CSPs and are affected by RNA binding. (E) C-terminal residues of Sxl-dRBD5 are not affected further upon addition of Unr-CSD1.

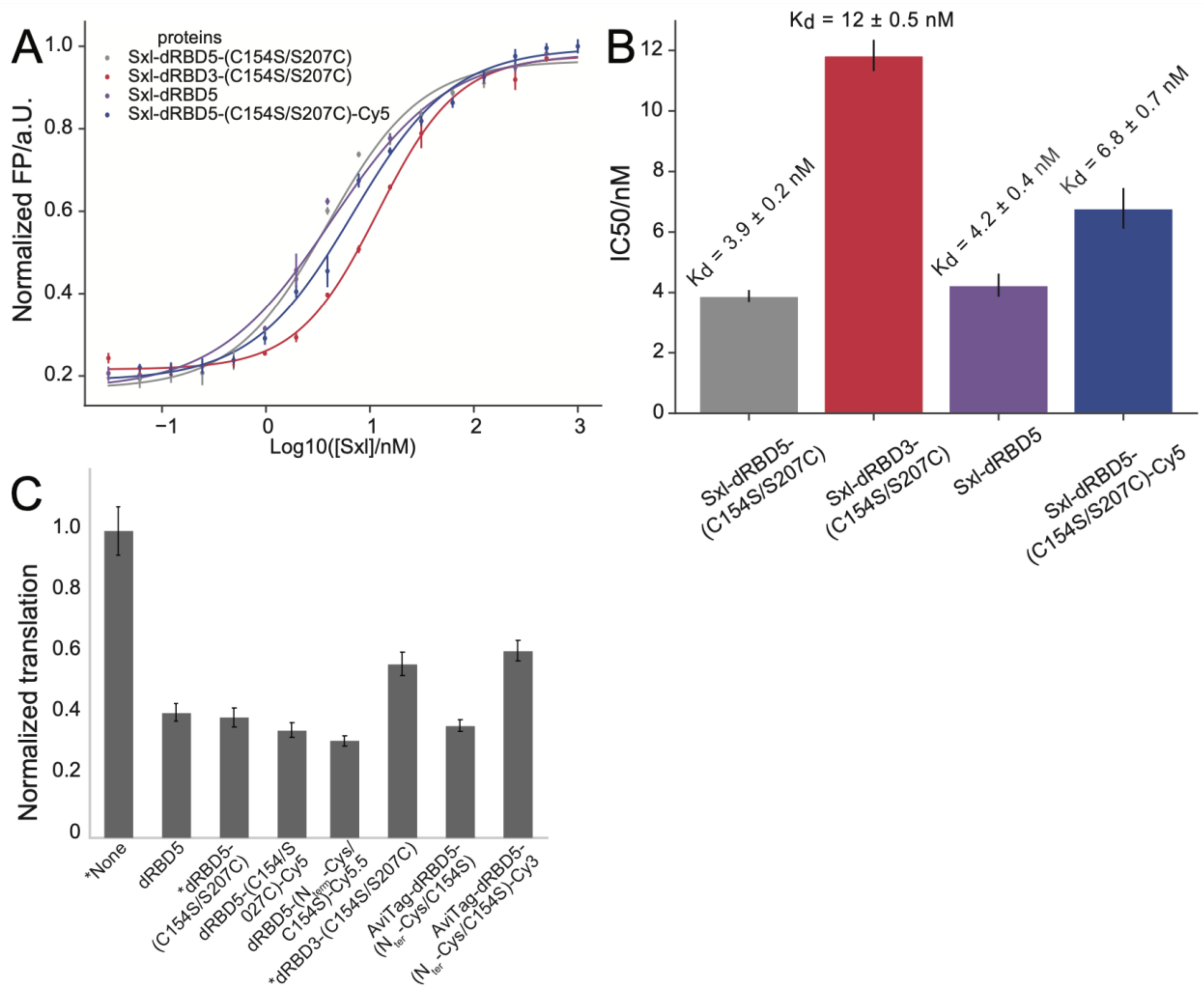


**Fig. S3. Affinity and activity of Sxl constructs, related to Fig. 2.** (A) Fluorescence polarization assay of different Sxl constructs to 3' FAM-labeled *msl-2*$_{Fb}$ mRNA to obtain (B) affinity values (IC$_{50}$) from fit curves. (C) Translation repression activity assay of a reporter mRNA containing a luciferase ORF and the minimal *msl-2* 3' UTR in the presence of various Sxl constructs (1.2 µM). Measurements were performed in triplicates. Conditions marked with asterisks are shown in Fig. 2D.

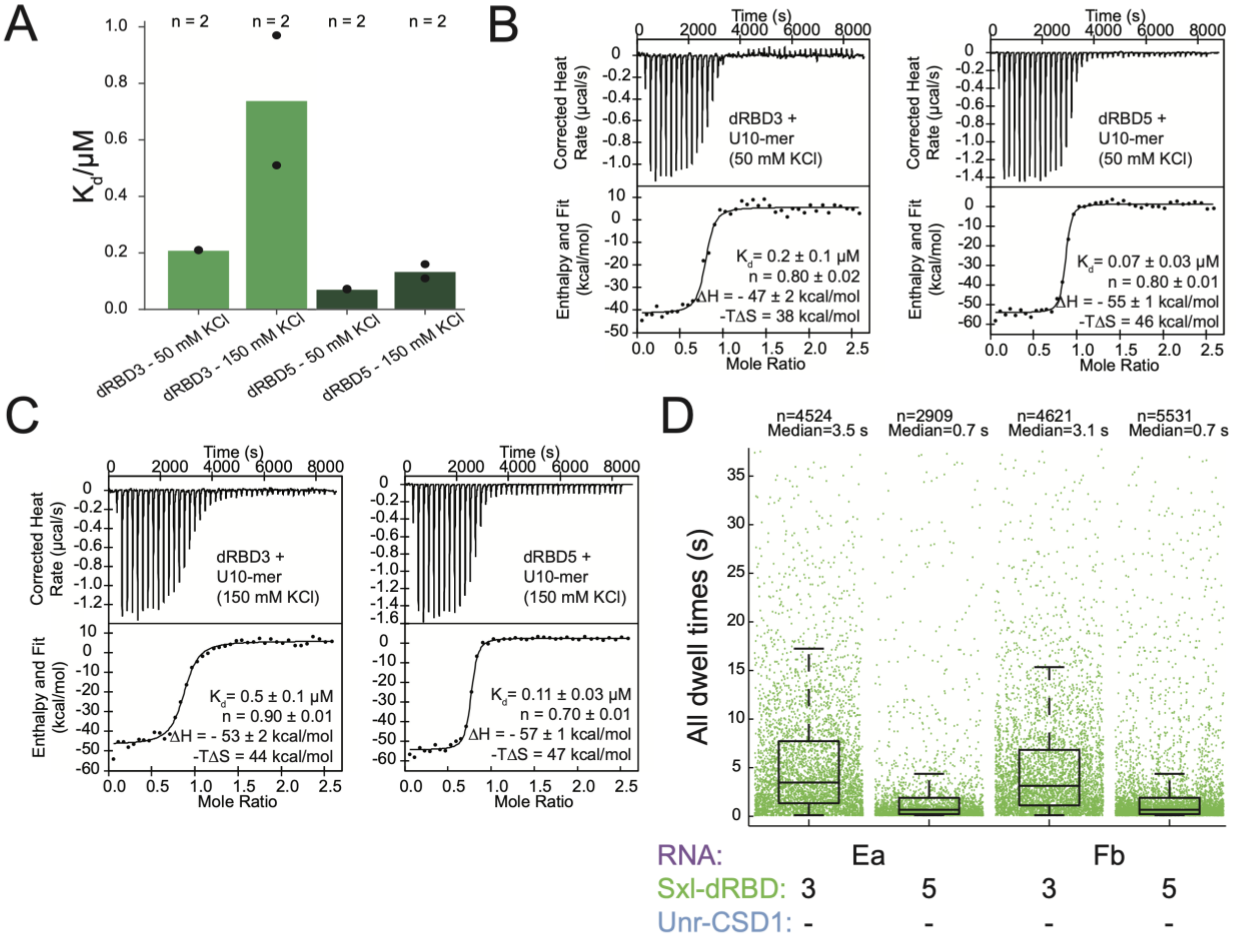


**Fig. S4. Affinity of Sxl constructs at different salt concentrations and arrival times of Sxl and Unr, related to Fig. 2.** (A-C) ITC measurements: (A) Overview of dissociation constants for Sxl-dRBD3 and dRBD5 are shown as replicates. Values were derived from fits of thermograms at 50 mM (B) and 150 mM (C) KCl, respectively. To avoid the effect of avidity on $K_d$ values, we determined binding of Sxl to a $U_{10}$-mer, the cognate binding motif of Sxl. (D) Arrival times determined from smFRET measurements. All arrival times were determined from Sxl-dRBD3 or Sxl-dRBD5 binding (40 nM total) to *msl-2*$_{Ea}$ and *msl-2*$_{Fb}$ mRNA.

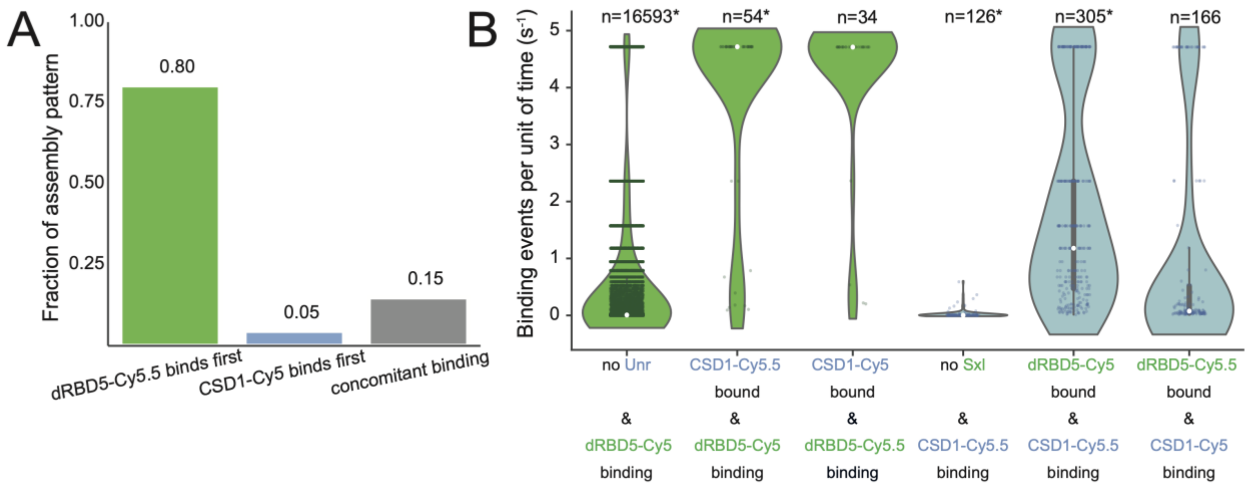


**Fig. S5. Multi-color smFRET assay kinetics, related to Fig. 3.** (A) The fraction of assembly patterns obtained in a multi-color smFRET assay in which *msl-2*$_{Fb}$ binding with Sxl-dRBD5$_{Cy5.5}$ and Unr-CSD1$_{Cy5}$ was probed. (B) A side-by-side comparison of the binding events per second for both labeling schemes (Sxl-dRBD5$_{Cy5}$ and Unr-CSD1$_{Cy5.5}$; Sxl-dRBD5$_{Cy5.5}$ and Unr-CSD1$_{Cy5}$) is shown. Violinplots marked with an asterisk are shown in Fig. 3D.

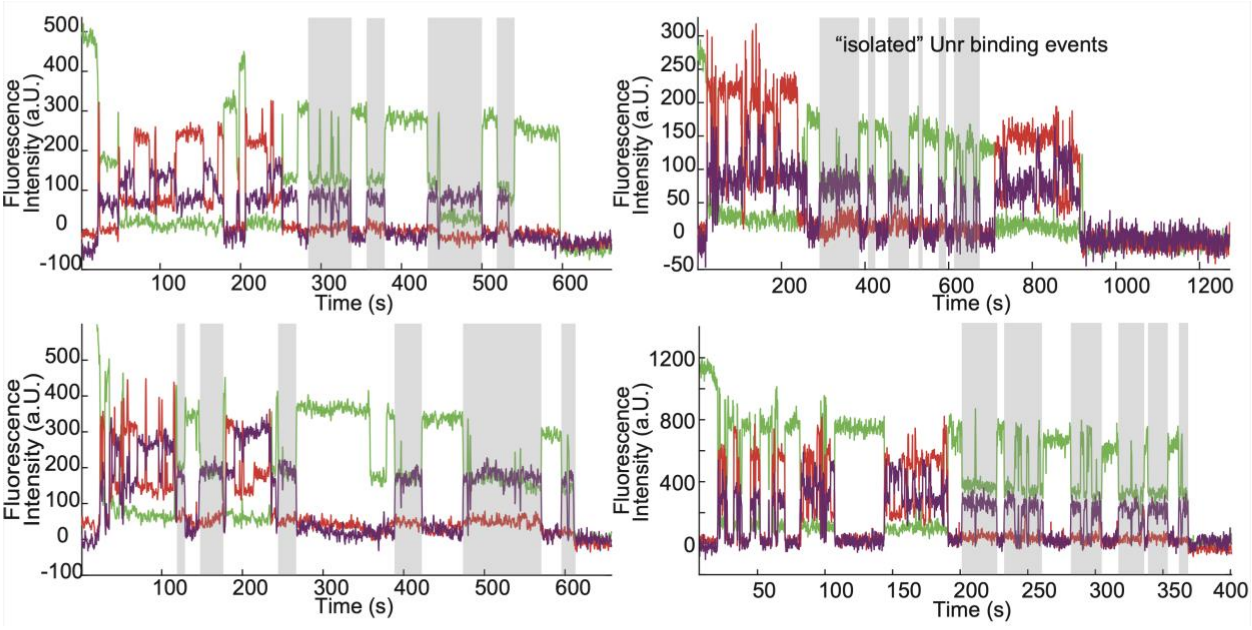


**Fig. S6. Multi-color smFRET assay, related to Fig. 3.** Exemplary traces of multi-color binding assays of Sxl-dRBD5-Cy5 (40 nM total) and Unr-CSD1-Cy5.5 (200 nM total) to *msl-2*$_{Fb}$. The long-lived “isolated” Unr-CSD1 binding events are highlighted in gray, which arise when unlabeled Sxl is co-bound (Sxl-dRBD5-Cy5 labeling efficiency is 70 %). The first part of the bottom right trace is shown in Fig. 3B.

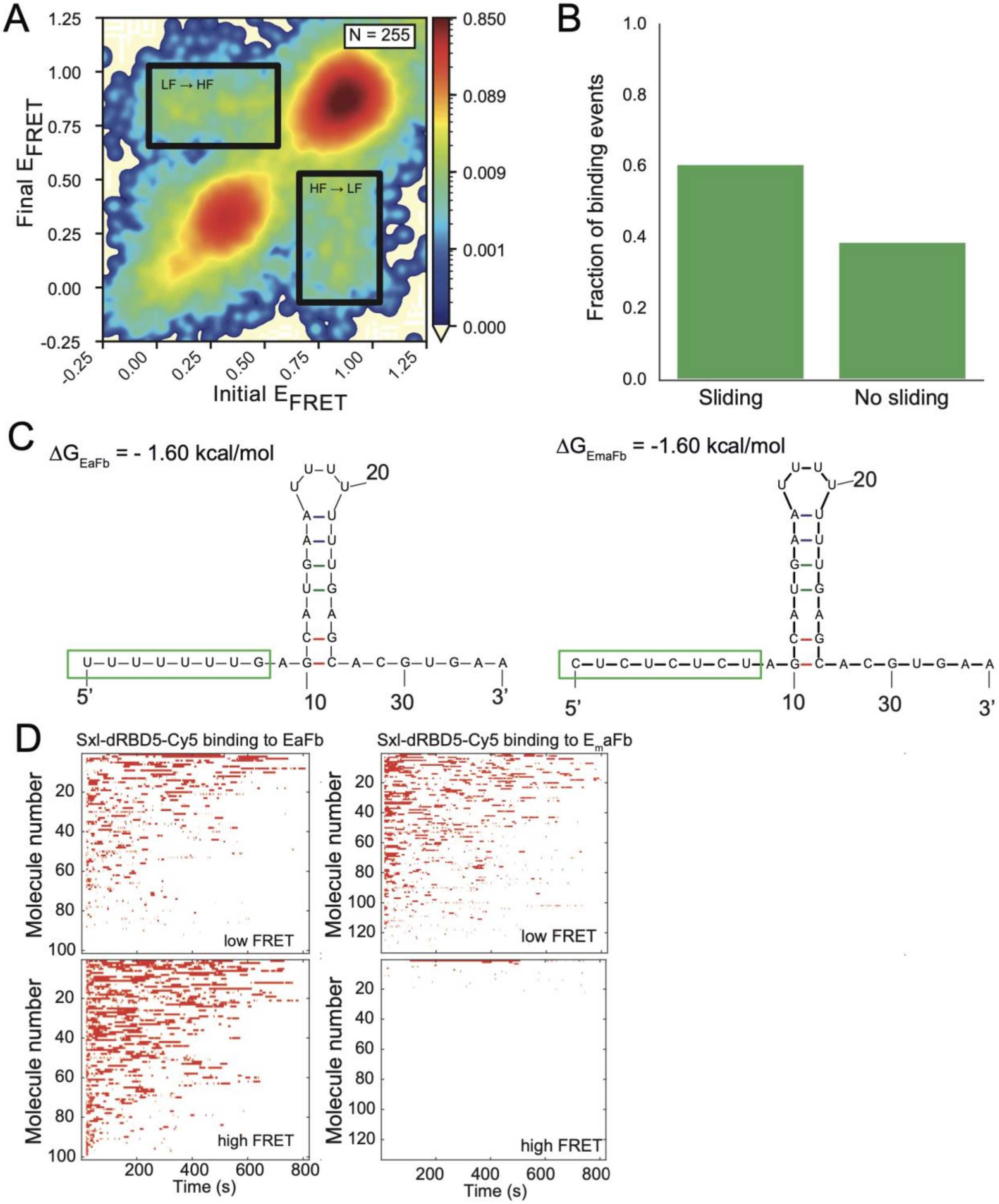


**Fig. S7. Sxl tandem RNA binding smFRET assays, related to Fig. 4.** (A) A transition density plot from Sxl-dRBD5-Cy5 binding events to immobilized *msl-2*$_{EaFb}$ shows transition from high to low FRET efficiency states and vice versa. (B) Fraction of binding events that exhibit Sxl sliding between the E and F sites. (C) Secondary structure predictions of EaFb and a Sxl binding site mutant (E$_m$aFb)(Zuker, 2003). (D) Clustered heatmaps (sorted according to total bound lifetime) of Sxl-dRBD5-Cy5 binding to different substrates shows that high FRET binding events are almost abolished in the Sxl E site mutant.

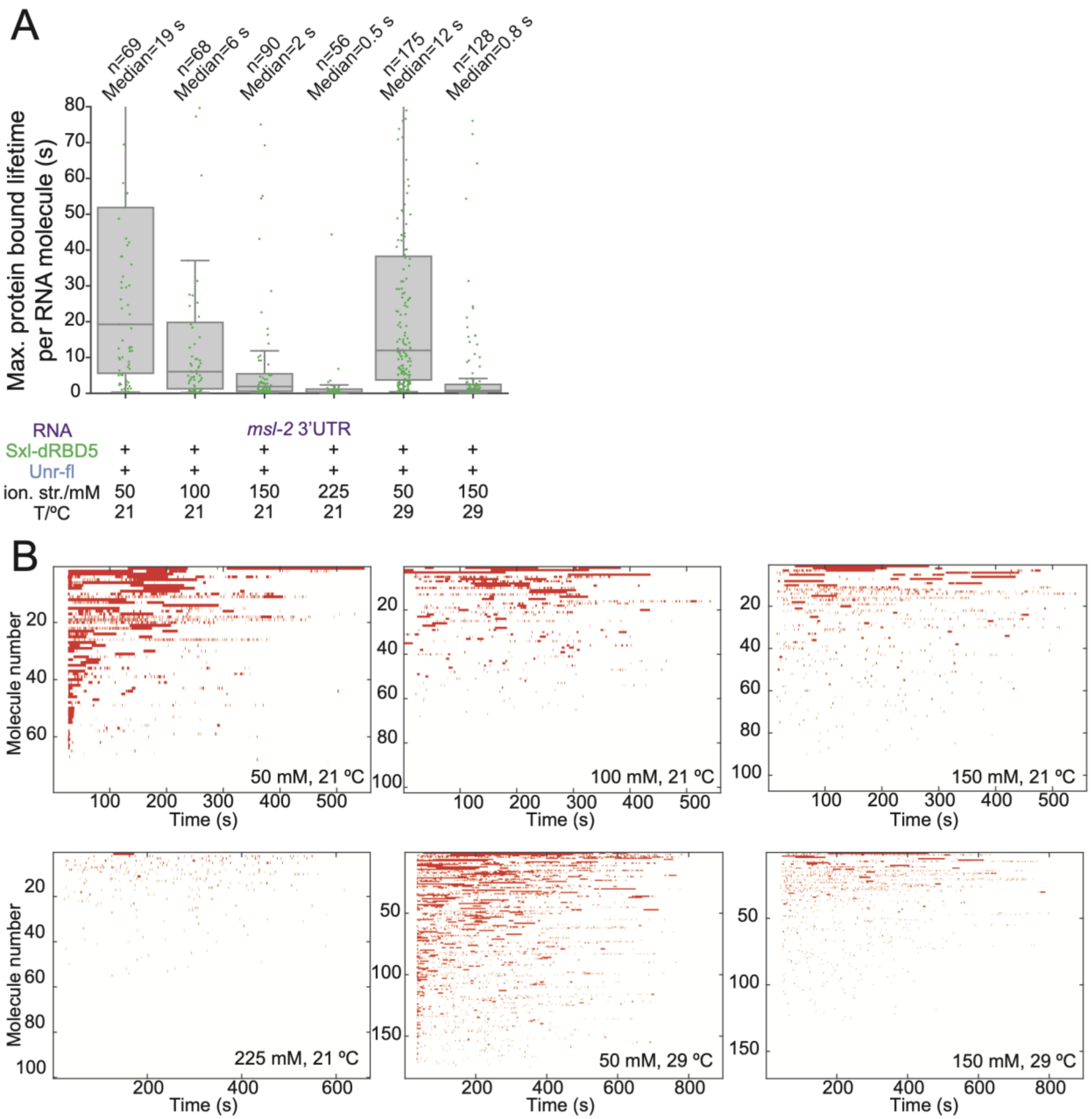


**Fig. S8. Sxl binding to *msl-2* 3' UTR at different ionic strengths and temperatures, related to Fig. 5.** (A) Maximum protein-bound dwell times of Sxl-dRBD5-Cy5 per *msl-2* 3' UTR molecule is shown for different ionic strengths (adjusted with NaCl) and different temperatures. (B) Clustered heatmaps sorted by total bound time at varying conditions.

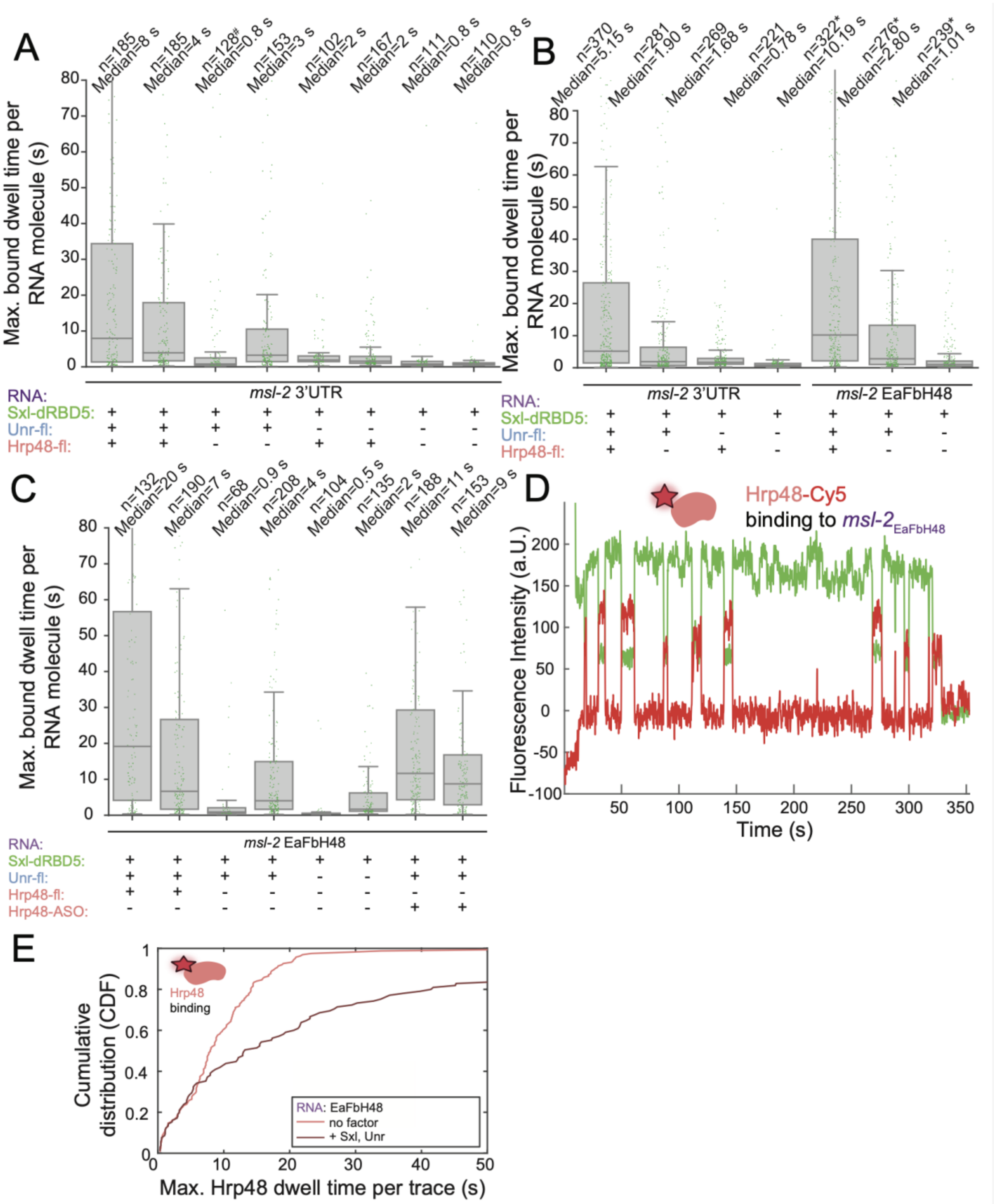


**Fig. S9. Hrp48 chaperoning activity, related to Fig. 5.** (A) A side-by-side comparison of duplicates. Differences in duplicates arise due to batch effects of individual Sxl and Unr purification batches. Sxl-dRBD5-Cy5 binding was tested in absence and presence of full-length Hrp48 and full-length Unr. Replicates marked with a hash are shown in Fig. S8A. (B) Merged duplicates (from (A) and (C)) of Sxl-dRBD5 binding to both *msl-2* 3' UTR and the shorter *msl-2*$_{EaFbH48}$ mRNA in presence and absence of other factors. Replicates marked with asterisks are shown in Fig. 5B. (C) Side-by-side comparison of duplicates measured for Sxl-dRBD5-Cy5 binding to a shorter RNA construct *msl-2*$_{EaFbH48}$. (D) Hrp48-Cy5 binding to *msl-2*$_{EaFbH48}$ in the absence of factors. (E) Cumulative distributions of Hrp48-Cy5 binding in absence and presence of Sxl-dRBD5 and full-length Unr.

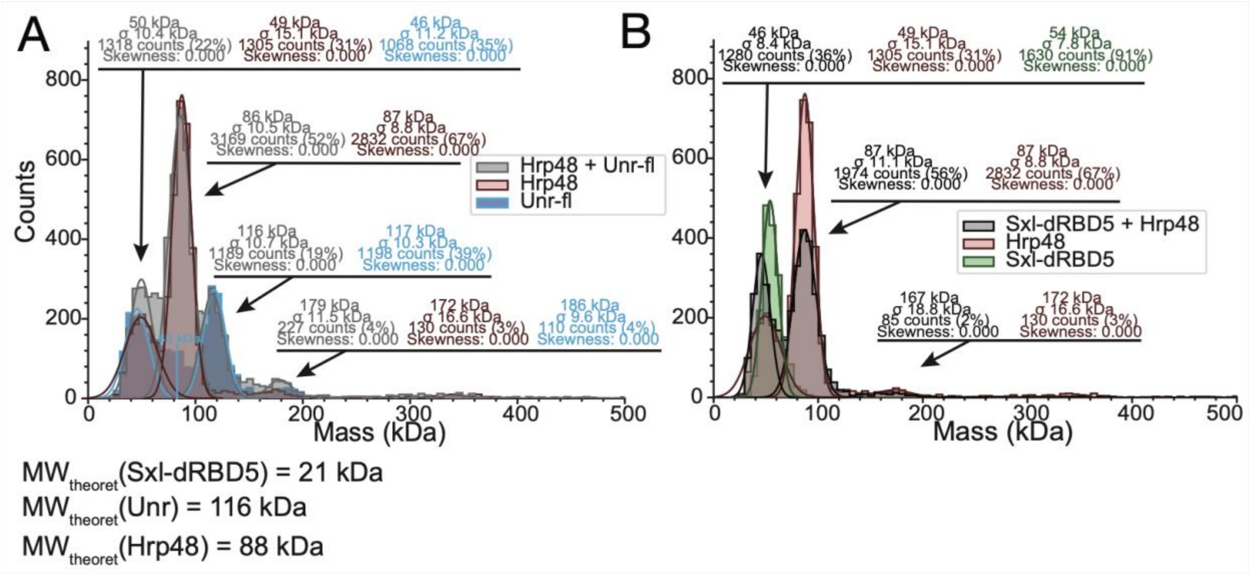


**Fig. S10. No complex formation of Hrp48 with Unr/Sxl observed as tested by mass photometry, related to Fig. 5.** (A) Probing interaction of Hrp48 and Unr. A small peak at ~ 180 kDa in Unr is due to a contamination and therefore does not indicate Hrp48 and Unr complex formation at an expected molecular weight of ~ 216 kDa. (B) Complex formation of Sxl and Hrp48 in the absence of RNA was also not observable with mass photometry.
